# Integration of allocentric and egocentric visual information in a convolutional / multilayer perceptron network model of goal-directed gaze shifts

**DOI:** 10.1101/2021.12.15.472867

**Authors:** Parisa Abedi Khoozani, Vishal Bharmauria, Adrian Schütz, Richard P. Wildes, J. Douglas Crawford

## Abstract

Allocentric (landmark-centered) and egocentric (eye-centered) visual codes are fundamental for spatial cognition, navigation, and goal-directed movement. Neuroimaging and neurophysiology suggest these codes are segregated initially, but then reintegrated in frontal cortex for movement control. We created and validated a theoretical framework for this process using physiologically constrained inputs and outputs. To implement a general framework, we integrated a Convolutional Neural Network (CNN) of the visual system with a Multilayer Perceptron (MLP) model of the sensorimotor transformation. The network was trained on a task where a landmark shifted relative to the saccade target. These visual parameters were input to the CNN, the CNN output and initial gaze position to the MLP, and a decoder transformed MLP output into saccade vectors. Decoded saccade output replicated idealized training sets with various allocentric weightings, and actual monkey data where the landmark shift had a partial influence (R^2^ = 0.8). Furthermore, MLP output units accurately simulated prefrontal response field shifts recorded from monkeys during the same paradigm. In summary, our model replicated both the general properties of the visuomotor transformations for gaze and specific experimental results obtained during allocentric-egocentric integration, suggesting it can provide a general framework for understanding these and other complex visuomotor behaviors.

## 1 Introduction

The visual system has two ways to code object location: relative to oneself (egocentric; Andersen & Buneo, 2002; Crawford et al., 2011) or relative to surrounding objects (allocentric; Schenk 2006; Ball et al. 2009; Chen et al. 2011). This distinction has proven to be fundamental in accounts of the role of the hippocampus in spatial memory (Rolls, 2019; Danjo, 2020) and in two-stream theories of vision (Schenk 2006;Thaler and Goodale, 2011). This is also true for goal-directed movements. When egocentric and allocentric locations conflict, participants can be explicitly instructed to use one or the other cue, but normally (when stable) they are integrated to minimize internal noise and thus reduce end-point variability in the behavior (Byrne et al. 2010; Chen et al. 2011; Li et al. 2017). Numerous behavioural studies have suggested that this involves a process similar to Bayesian Integration (Neggers et al. 2005; Byrne & Crawford 2010; Fiehler et al. 2014; Li et al. 2017; Klinghammer et al. 2017; Klinghammer et al. 2018). However, the intrinsic mechanisms for representing and integrating these codes remains a puzzle.

Cue-conflict behavioural studies combined with theoretical modeling were used first to investigate the computational rules for allocentric and egocentric integration for reach control (Byrne and Crawford 2010; Fiehler et al. 2014; Klinghammer et al. 2017; Klinghammer et al. 2018). For example, when a landmark is shifted relative to the remembered location of a reach target, reach end points shift partially in the same direction, consistent with Bayesian integration and the outputs of a maximum likelihood estimator (Byrne and Crawford 2010). The amount of the shift was affected by the number of relevant objects in the scene as well as scene consistency (Klinghammer et al. 2017, 2018). Neuropsychology results suggest that egocentric and allocentric visual representations are segregated in the dorsal and ventral streams of vision respectively (Schenk, 2006; Thaler and Goodale, 2011), suggesting a need to reintegrate this information at some point in the brain. Subsequent neuroimaging studies confirmed such segregation (Chen et al. 2014) and suggested that the recombination occurs in parietofrontal cortex (Chen et al., 2018), but neuroimaging data are too coarse-grained to identify specific computational mechanisms at the cell / circuit level.

These observations also have been extended to the monkey gaze control system. As in reaching, when monkeys were trained to saccade to remembered targets in the presence of a landmark shift, their gaze endpoints shifted partially in the same direction, again suggesting weighted integration between egocentric and allocentric inputs (Li et al., 2017). Landmark influence was larger when it was closer to initial gaze fixation and when it was shifted away from the target. The development of this animal model then allowed for neural recordings associated with allocentric-egocentric integration for goal directed movements. In particular, Bharmauria et al. (2020, 2021) recorded from the frontal eye field (FEF) and supplementary eye fields (SEF) while rhesus monkeys performed the cue-conflict gaze task described above. When the landmark shifted, its influence was first observed multiplexed in FEF/SEF delay responses, and then fully integrated in the gaze motor response (Bharmauria et al., 2020, 2021).

In summary, it appears that egocentric and allocentric codes are at least partially segregated within the visual system, but then reintegrated for action in parietofrontal cortex. However, the early cellular / network mechanisms for these processes remain unknown. One reason for this knowledge gap is the lack of a theoretical framework to guide physiological studies in this area. Specifically, there is a lack of models that integrate both the complexity of the visual system and the sensorimotor system in general, and allocentric-egocentric integration in particular.

One approach to building such a theoretical framework is to deploy and analyze artificial neural networks (ANNs). But here again is a knowledge gap because this phenomenon requires the integration of multiple features into the final motor output. Many ANN models have attempted to represent feature interactions in the visual system, but do not consider sensorimotor transformations (e.g., Geirhos et al., 2017; Rajalingham et al., 2018; Schrimpf et al., 2018; Kar et al., 2019). Conversely, models of goal-directed behavior typically focus on the sensorimotor transformation and treat the visual world as a single ‘dot’ (Zipser & Anderson, 1988; Smith & Crawford, 2005; Blohm et al., 2009). What is needed, is a more general model with the capacity to represent both multiple object features and implement the sensorimotor transformation.

The purpose of this research was to develop and validate a neural network framework for representation and integration of allocentric and egocentric information in the visuomotor system, with the ultimate goals of understanding the current data and generating predictions for new experiments. To achieve these goals, we used a Convolutional Neural Network (CNN) to represent the visual system, connected in series with a Multi-Layer Perceptron (MLP) network to represent the visuomotor transformation. We modeled saccades because of their simplicity and the availability of relevant data. To ensure physiological relevance, we employed fully analytic solutions (Hadji & Wildes, 2017), with inputs based on known properties of the sensory cortex (Hubel & Wiesel, 1968; Carandini et al., 2005; Carandini, 2006; Wang et al., 2007) and outputs based on the motor response field properties of the frontal eye field (Knight & Fuchs, 2007; Knight, 2012; Sajad et al., 2015; Caruso et al., 2018). To train this network, we used both synthetic datasets and actual data obtained from the cue conflict task described above (Li et al., 2017; Bharmauria et al., 2020, 2021). Finally, we compared the network properties and outputs to this data. The results suggest that our network captures the known behavioral and neural properties of sensorimotor system, and thus provides a promising framework for understanding and predicting unknown properties. Further, the model has potential for application in a wide range of scenarios involving the integration of visual features for complex visuomotor tasks.

## 2 Method

To create our model, four main challenges were addressed. First, training an ANN with images requires large datasets (Goodfellow et al. 2016), which is infeasible to create considering the complexity of the experiments employed in this field. Second, to yield results that are physiologically relevant, it is essential to incorporate the known physiology into the ANN. This is particularly challenging to solve for the current question because it requires modeling both the early visual system and the visuomotor transformation for goal-directed saccades, a task that (to our knowledge) has not been attempted before. Third, it is desirable for the internal operations of the model to be interpretable, which is a lack in typical learned ANNS. Finally, it was necessary to train and validate our model against real data. To do this, we used a recent experiment in our lab to train our network (Li et al. 2010; Bharmauria et al. 2020, 2021). Therefore, in the following, we first explain the task on which we based our simulation. Then, we explain the model in a more general format. Finally, we explain the extracted data we used for our simulations and the model parameters.

### 2.1 Task

The details of our model were motivated based on the insights from recent experiments in the macaque gaze system (Li et al. 2010; Bharmauria et al. 2020, 2021). Therefore, it is essential to first summarize the task (Figure 1A). Initially, a red dot appears on a screen and monkeys are trained to fixate their gaze on the red dot. After the fixation, an image, which consists of allocentric cues (two crossing lines) and a visual target (white dot), appears on the screen (encoding image), while the fixation point remains. Monkeys have 100ms to memorize the position of the target. After a delay, a mask appears and after a further random delay period, a decoding image appears. The decoding image consists of only the landmark. Based on the experiment protocol, the landmark location can be the same as it was in the encoding image (no shift condition) or it can be different (shift condition). The task for the monkey is to saccade toward the missing target location when the fixation signal (red dot) disappears. Monkeys were rewarded if their gaze landed within 8°-12° of the target. The landmark locations were randomly selected from four possible locations distributed on the edges of a square and 11° away from the target. The shift direction was randomly selected amongst 8 uniformly distributed points on a circle 8° away from the initial landmark location. Additionally, the initial gaze location (fixation) was jittered within a 7-12° window (Figure 1B). As noted in the introduction, the results showed an influence of the landmark shift on gaze behavior as well as on SEF/FEF motor neuron population code and intrinsic coordinate frames.

**Figure 1.**
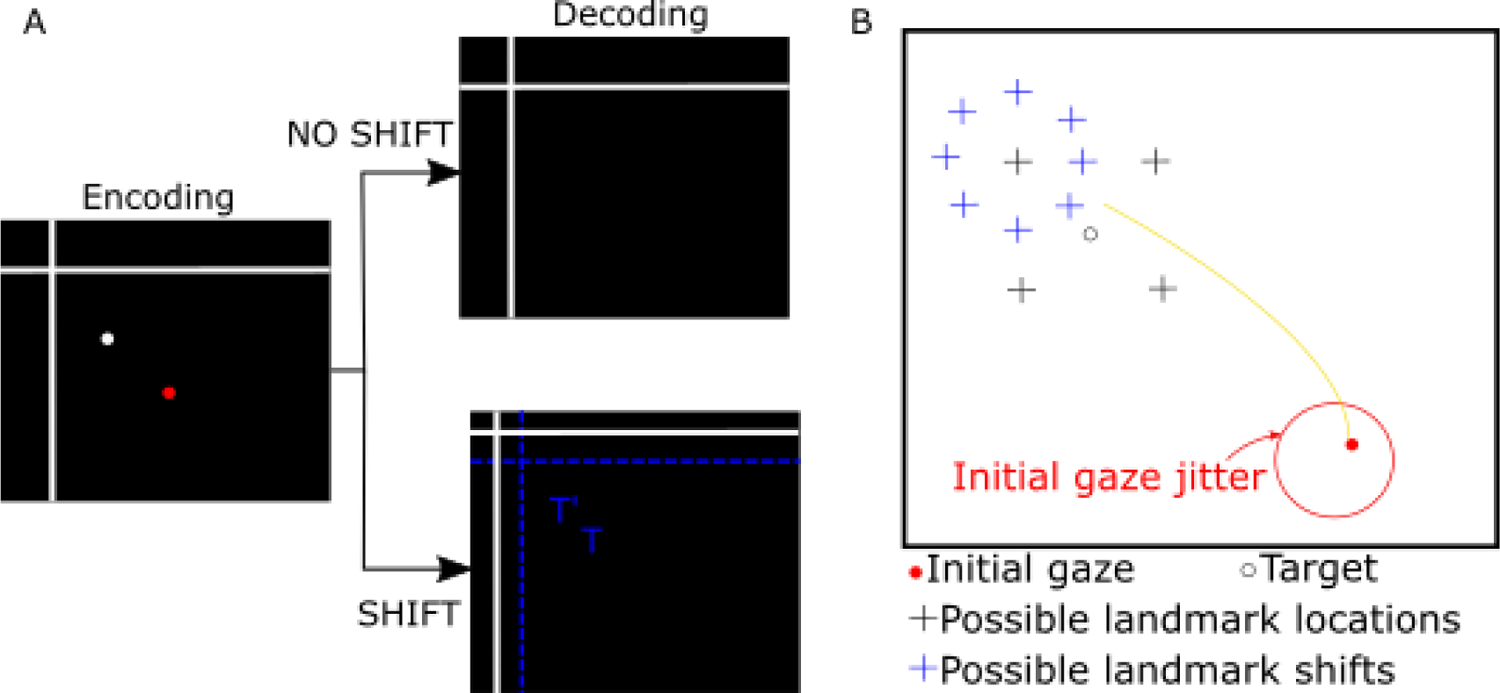
Experimental conditions. **A)** After initial fixation on a red dot, an image consisting of allocentric cues (two crossing lines) and visual target (white dot) appears on the screen along with the fixation dot. We consider this image an encoding image. Monkeys have a limited time to memorize the position of the target from this image. After a random delay during which the encoding image is absent, the decoding image consisting of only visual landmarks and fixation dot appears. Based on the experiment protocol, the landmark location can be the same as the encoding image (no shift condition) or can be different from the encoding image (shift condition). The task for the monkey is to perform a saccade toward the missing target location when the fixation signal (red dot) disappears. B) Landmark location (black crosses) was randomly selected from 4 possible positions, 11° from the target (open circle). Landmark shift was randomly selected from eight possible shifts (blue crosses) evenly distributed on a circle 8° from the initial landmark location. Initial gaze location was jittered within a 7-12° window. Illustrated shift positions are with respect to the landmark at their centre; shifts for other depicted landmarks not shown for sake of simplicity of illustration.

### 2.2 Theoretical model

#### 2.2.1 Model overview

Our theoretical model provides a framework to study the combination of allocentric and egocentric information for goal-directed reaching movements in the brain. Toward this end, we combined two current approaches in machine learning (Goodfellow et al. 2016): a Convolutional Neural Network (CNN) and a Multi-Layer Perceptron (MLP) to simulate different modules of this transformation in the cerebral cortex. In this section, we provide an overview of the proposed network and the input-output structure. Note that since inputs and outputs of the model represent two-dimensional (2D) directions in eye coordinates, we have simplified the model by implementing it in 2D.

Our proposed network comprises four main stages (Figure 2A): inputs, a convolutional neural network, a multilayer perceptron, and output. The inputs to the network provide necessary information for the combination of allocentric and egocentric information. We feed the network with images that consist of visual targets and surrounding landmark cues, based on recent studies in the macaque, as described in the previous subsection, to provide allocentric information. These images are generated in spatial coordinates and transformed into eye-coordinates to mimic retinal projections (e.g., Klier & Crawford 1998; Klier et al. 2001). Since we did not require this model to perform 3D transformations, the only extraretinal signal that we provided was a 2D gaze signal. The second stage of our network is a CNN (Figure 2B), which was deployed to create abstract representations of input images. The challenge here was to design the CNN as a physiologically plausible model of the early visual cortex (Hubel & Wiesel, 1968; Carandini et al., 2005; Carandini, 2006). In the third stage, we used an MLP (Figure 2C) to perform the required sensorimotor transformations. The fourth stage, the output, provides the decoded saccade vectors (e.g., displacement of the eye, coded as final gaze position in eye coordinates) from motor populations. Details of the implementation of these four network stages (in the format of input – output and hidden units: CNN and MLP) are provided in subsequent sections.

**Figure 2.**
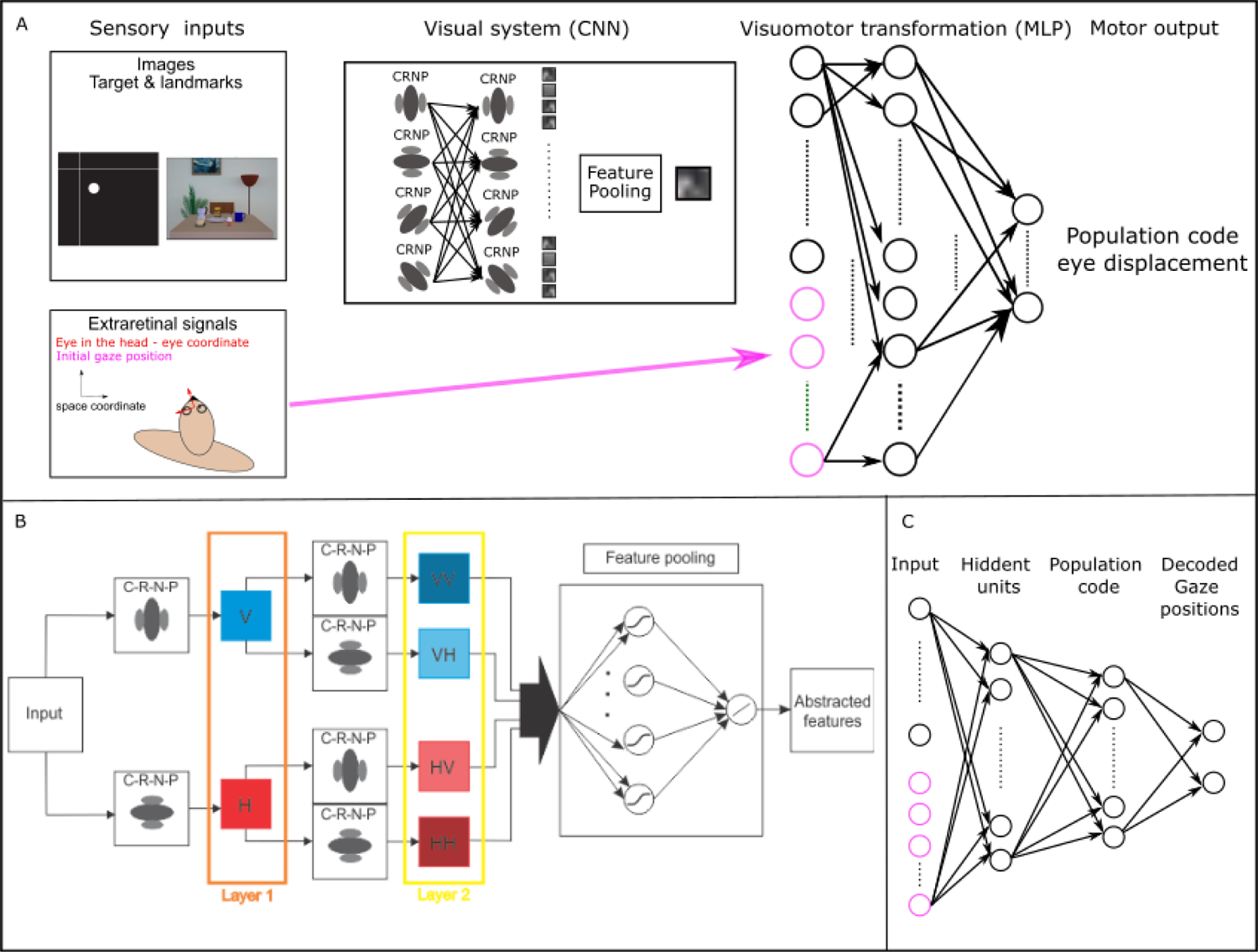
Proposed model. **A)** Overall network architecture. Our network consists of 4 main components. 1) input signals: retinal images that contain visual targets and allocentric landmarks, and extra-retinal signals i.e., eye position. We used 2D eye position for our simulations, 2) a convolutional neural network to create an abstract representation of the retinal images, 3) a multi-layer perceptron to implement the required sensorimotor transformations, and 4) output signal: a decoder to transform the population code of the required movements (eye displacement) into a saccade vector. B) CNN network architecture and *Repeated filter mechanism.* Layer 1 extracts local spatial features. C-R-N-P represents convolution, rectification, normalization, and spatial pooling. Additionally, V and H represent Vertical and Horizontal filtered results, respectively. A network with only two layers and two sets of filters is depicted for illustrative purposes; four filters are used in actual implementation, i.e., adding two more oriented along the diagonals. Each feature map at Layer 2 is treated as a new separate signal and is fed to the feature pooling layer. Symbol strings (e.g., VV, VH, etc.) indicate repeated filtering. The feature pooling layer consists of two fully connected layers: the first layer with a sigmoid transfer function and the second layer with a linear transfer function. The feature pooling layer combines the feature map from the final filtering layer to create an abstract representation of the required features. C) *MLP network architecture.* Our MLP network is a fully connected feed-forward neural network with four layers. The first layer is the input layer comprised of the extracted features from the images as well as extra-retinal signals. The second layer is comprised of our hidden units with sigmoid transfer functions. The third layer is the population code of the required motor movement for reaching toward the visual target. Finally, we added a read-out layer with fixed connection weights to decode the population code into 2D reach movements in Euclidean space.

#### 2.2.2 Model Input-output signals

The network required two types of input signals: 1) retinal images specifying allocentric landmarks and the visual target position (which provides allocentric information); 2) extra-retinal signals (which provide egocentric information), which in real-world conditions are required for accurate coordinate transformations (Crawford et al. 2010) and calculation of motor error (ME). The output is the signal that drives the effector (e.g., the eye in this case) from its current position to the target position (ME coded in a population format). Here, we modeled/simulated a simplified 2D representation of the retina and gaze to focus on the issue of allocentric-egocentric integration.

##### 2.2.2.1 Visual inputs

The first component of the input for the network is the simulated retinal image. This component simulates a projection of the world containing both the movement target and additional landmarks on the retina. These inputs corresponded to the encoding and decoding images of the task defined in Section 2.1. Since we are interested in investigating the spatial component of the allocentric and egocentric combination, and save modeling of temporal aspects for future work, we process the two images separately as input to our network, each image having dimensions WxH, with the W and H width and height, respectively.

To simulate these inputs, we generated two sets of images: encoding and decoding. Each image, with the size of 200×200 pixels, contains two crossing lines (a horizontal line and a vertical line, each of width 1 pixel and length extending across the entire image) as well as a 6×6 pixel square representing the visual target. The image intensity values for the target and lines was set to 1 and the rest of the images were set to 0. Based on the experiment protocol, there were two conditions for decoding images: The landmark appeared in the same location as the encoding image; or the landmark appeared in a shifted location compared to the encoding image. The specific values for the landmark and target positions for both encoding and decoding images were extracted from the actual experiment’s protocol (for further details refer to Bharmauria et al., 2020). Retinal images were calculated based on the spatial configurations of the stimuli relative to initial eye (gaze) positions. To perform this calculation, we deployed a simplified 2D equation (torsional factor is ignored for now) and calculated the retinal image by subtracting the eye position from the actual images. These retinal image values were then used as inputs the first layer of our convolutional network, described below.

##### 2.2.2.2 Eye position signal

As mentioned above, extra-retinal signals are essential for performing the required reference frame transformations for various sensorimotor behaviors (Soechting & Flanders, 1992; Andersen & Buneo, 2002; Crawford et al., 2011). Although this transformation was not a focus here, we included a minimal extraretinal input (initial 2D gaze position) so that the model has the potential to be generalized to other situations. We used the angle vector representation, (r_x_, r_y_, r_z_), which represents the unit vector multiplied by rotation in degrees. To encode the angular value along each axis, we used a coding mechanism analogous to activities reported in the somatosensory cortex for coding eye position (Wang et al. 2007): 1) Neurons have Gaussian receptive fields with the peak indicating the preferred gaze direction; 2) the preferred direction of neurons does not form a topographic map; 3) a neuron’s activity at the peak of the receptive field is monotonically increasing as the saccade amplitude grows. Similarly, we used Gaussian receptive fields randomly distributed around the orbit and gain modulated by the saccade amplitude to code the eye position in our network. This coding mechanism is formulated as:

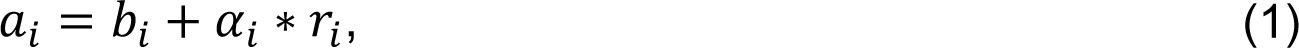

where *a_i_, b_i_, α_i_*, and *r_i_* represent the *i^th^* unit’s activation, baseline activity, rate of the activity increase, and angular direction, respectively. Baseline activity and the rate of growth in activity are randomly picked from (0, *max_bi/αi_*). A similar mechanism is used for coding hand or head position (King et al., 1981; Fukushima et al., 1990; Xing & Andersen, 2000). Therefore, our model can be extended to include additional extra-retinal signals, if needed.

##### 2.2.2.3 Motor output signal

The goal of the network is to generate movements toward a visual target. Previous work (Bruce & Goldberg, 1985; Sommer & Wurtz, 2006; Sajad et al. 2015, 2016) showed that the motor neurons in frontal eye field represent an open-end receptive field: The neural activity increases with increasing saccade amplitude. In contrast, theoretical studies suggested that cosine tuning is optimal for motor control in 3D (Kalaska, et al., 1997; Scott, 2001; Kakei et al., 2001, 2003). To reconcile the results of these studies with our data, we represented the response fields of our output layer as the first quarter (0-90°) of a cosine curve, according to:

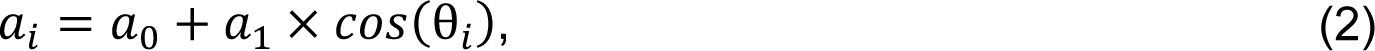

where *a_0_* = 0.5 is the baseline activity, 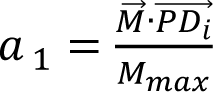 is the scaling factor, angle 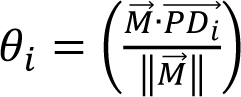 is the direction of the movement, *M_max_* is the maximum amplitude of the movement, 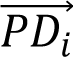 is the required movement, and 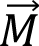 is the preferred movement direction of the unit. We use four or 100 units to represent motor neurons. We used 4 units for behavioural analysis and 100 units for neural analysis. The maximum movement range is 150 cm. Notably, in the actual data the influence of the landmark shift was fully integrated into the frontal eye field motor output signal, which coded a landmark-shifted gaze position in eye-centered coordinates (Bharmauria et al. 2020).

We also include an extra layer (i.e., linear decoder) to our network, called read-out, with two units: One coded the horizontal component of the movement, the other coded the vertical component. It has been shown that a linear decoding is suited specifically for arrays of neurons with tuning curves that resemble cosine functions (Salinas & Abbott, 1994). Therefore, we determined the required weights between the population code and read-out layer by an Optimal Linear Estimator (OLE; Salinas & Abbott, 1994),

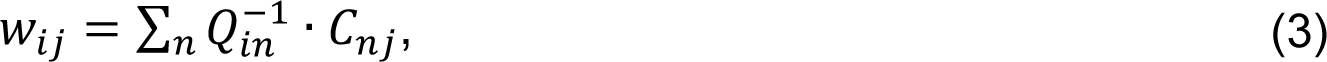

where *W_ij_* is the weight between unit *i* at population code to unit *j* at the read-out layer. Since we used a 2D representation, *j* is equal to 2 and consequently *M_1_* represents the horizontal value of movement and *M_2_* represents the vertical value of movement. Additionally, *Q_in_*, the correlation between firing rates of neuron *i* and *n*, and *C_nj_*, the center of the mass for the tuning curve function for the neuron *n*, are calculated as

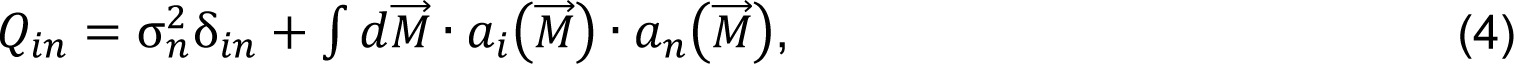

and

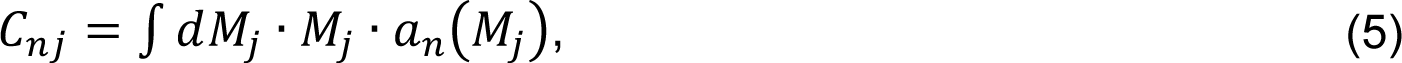

where *σ^2^_n_* is expected neural noise. For detailed discussion and calculations, see Salinas & Abbott (1994) and Blohm et al. (2009). These weights are fixed in the network and are not changed during training. Using this strategy, we enforced a more physiologically feasible population code for our network, which enabled us to have an unambiguous interpretation of single-unit activity. Additionally, using a read-out layer enabled us to have a more stable fitting procedure for training our network. Previous studies deployed a similar strategy (e.g., Smith and Crawford 2005; Blohm et al., 2009; Keith et al., 2010).

#### 2.2.3 Hidden units

We used two types of networks as our hidden units. The logic is to provide a similar paradigm as the brain, with one (CNN) representing the early visual areas detecting the relevant visual features and the other (MLP) representing the higher cortical layers performing the required computations (here sensorimotor transformations) for planning the appropriate action. The parameters of our CNN were predetermined based on analytic and physiological considerations, except the feature pooling layer, which was learned. The parameters of the MLP were determined based on the training using different datasets.

#### 2.2.4 Convolutional neural network (CNN)

In recent years, convolutional neural networks have shown promising performance for a wide variety of computer vision tasks such as image classification (for a review see, e.g., Liu et al., 2020). Some studies took a step further and showed that the learned filters in convolutional networks replicate neurons’ behaviour at the early visual cortex (e.g., Kriegeskorte, 2015). In general, a convolutional network is comprised of several main components: convolution, rectification, normalization, and pooling layers, which typically are repeated over several layers. The first challenge in designing a CNN is to set the required components and their associated parameters properly. Here, we briefly explain the rationale behind choosing our network components and parameters based on recent physiological findings.

One of the main challenges in processing visual scenes is extracting useful information from input images (e.g., spatial and spatiotemporal patterns indicative of objects, texture and motion). Hierarchical representations provide a powerful tool to address this challenge by progressively obtaining more abstract features at each successive layer. This incremental formation of more abstracted features yields a powerful data representation. A similar procedure is observed in the human brain. In the early visual cortex, a cascade of simple and complex cells is suggested to play a crucial role in extracting first-order stimuli in visual scenes (e.g., local measures of orientation; Hubel & Wisel, 1962; Heeger, 1991).

Additionally, a model with “filter-rectification-filter” has been suggested to be responsible for extracting higher-order patterns and creating a more global and abstract data representation in biological systems, e.g., texture (Baker & Mareschal, 2001). Likewise, it has been shown that a similar analogy of repeated filtering and rectification can generate state-of-the-art results in computer vision applications such as dynamic texture recognition (Hadji & Wildes, 2017). Here, we propose to use the same approach: repeated filtering. Figure 2B provides an overview of the convolutional network architecture. We only show a limited number of layers for illustrative purposes and only two filters in this figure. As mentioned above, a cascade of simple and complex cells is observed in the early visual cortex. Analogously, we employ a series of convolution, rectification, normalization, and pooling (C-R-N-P) processes to mimic the simple and complex cells’ functionality. The output of these C-R-N-P processes is then passed through the same procedures to yield the repeated filtering approach. At the last layer, we propose a cross-channel feature pooling process to create the final abstract representation of the data.

Figure 3 illustrates the efficacy of the repeated filtering approach. Figures represent the local energy in images at each layer. The input is an image that contains a vertical and a horizontal line crossing and a target represented as a dot, analogous to the stimuli used in Bharmauria (2020). The goal is to capture the crossing point and the target position. Here, we represent only two filter orientations: Horizontal (H) and Vertical (V). Three properties emerge after passing the image through the first layer filters: 1) the horizontal and vertical line are separated, 2) a gap is present at the mid-point of the line because of interference from the orthogonal lines in that vicinity, and 3) the target appears in both filterings, because its local shape drives both. Although the target is present in both filterings, its properties are now different, e.g., the target is mainly represented by vertical components after passing through the vertical filter. In the second layer, the distinction between the two-crossing lines becomes almost absolute. For instance, passing the input image through two vertically oriented filters (i.e., VV) resulted in the removal of the horizontal line. Also, the gap in the line is lessened because the interfering horizontal was suppressed from the previous layer of filtering. On the other hand, despite its different activation levels, the target is still present in all the filtered images. This result property enabled us to create an abstract representation of the target and landmark locations from input images using a feature pooling layer. In particular, our feature pooling layer detects common activated areas in all the filtered images at the second layer. Using preliminary experimentation, we found that two layers of repeated filtering is sufficient four our models. However, as shown in Figure 3 (and explained theoretically elsewhere, Hadji & Wildes 2017), the energy in the images decreases after each layer of filtering and consequent processes, and thereby has potential to provide an automatic criterion for determining the appropriate number of layers. In the next four sections, we detail the operation of each of the convolutional, rectification, normalization, and pooling layers.

**Figure 3.**
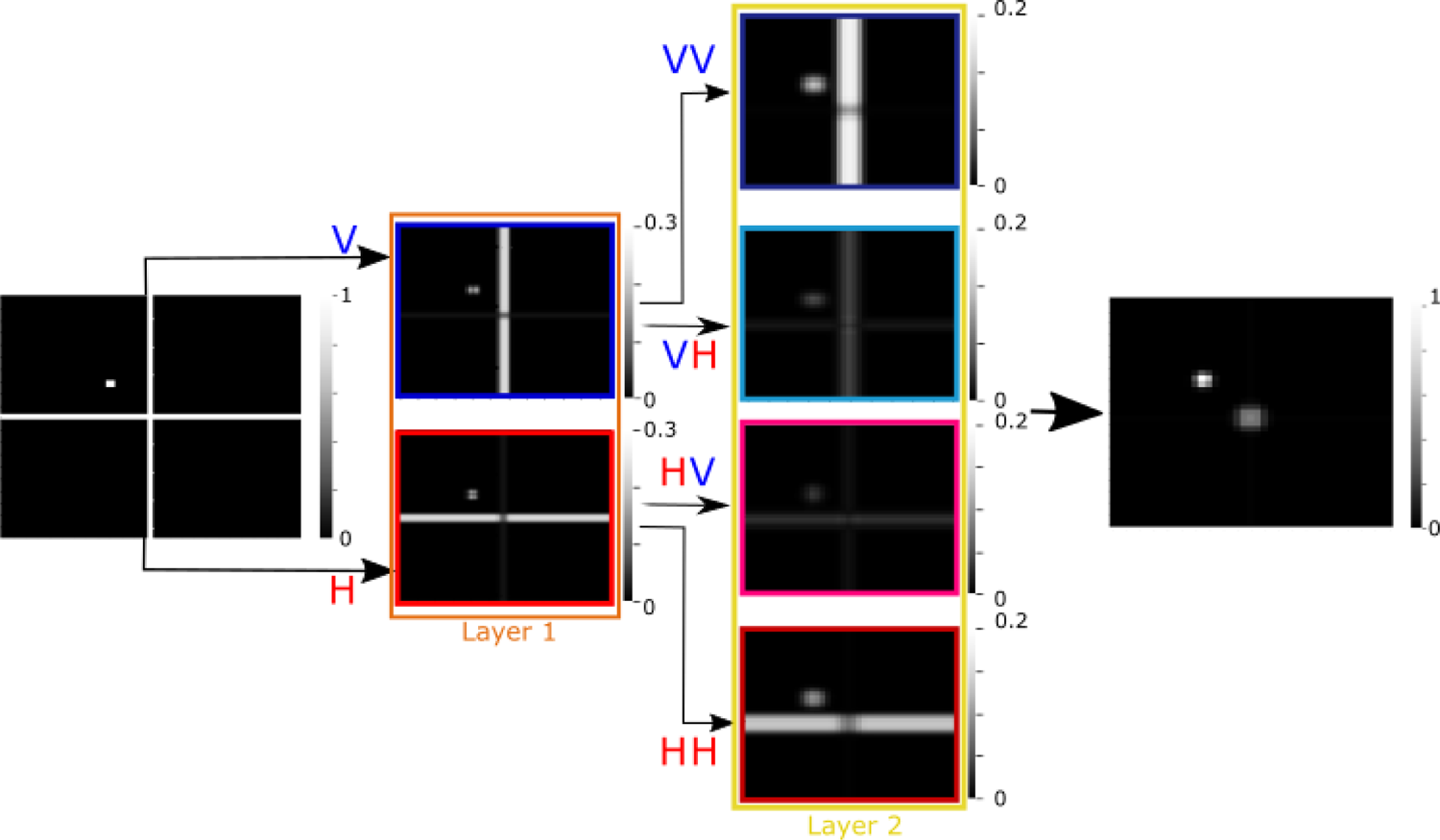
An example of processed images in a sample network with two layers and two sets of filters. V and H represent vertical and horizontal, respectively. Passing the input image through horizontal and vertical filters resulted in a separation of the horizontal and vertical lines. The target is present in both filtered images but now with different properties: more vertically oriented after passing through the vertical filter and more horizontally oriented after passing through the horizonal filter. Passing through the same filter twice (e.g., VV) resulted in almost full removal of the orthogonal line (e.g., horizonal line in VV). However, the target is still present at all the filter images with different activation level. Finally, combining all the locally extracted features by the trained feature pooling layer enabled us to detect the required features (crossing point and target position). Filter responses are represented as image brightness, with greater brightness corresponding to larger response.

##### 2.2.4.1 Convolution

The convolutional layer is one of the essential components of CNNs. Convolution is a linear, shift-invariant operation that performs a local weighting combination (filtering) across the input signals. The strength of such a function is that it extracts different input signal features based on the combined weights. Inspired by neuronal receptive fields in the early visual cortex, a set of oriented Gabor filters (Gabor, 1946) is a popular choice (Carandini et al., 2005; Carandini, 2006), which we symbolize as

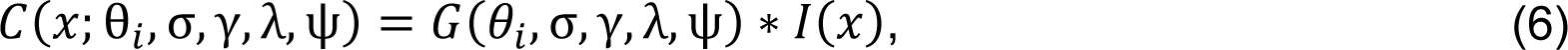

where *I(x)* is the input image parameterized by spatial position, *x, θ_i_* is the orientation of the Gabor filter *G*, while *σ, γ, λ, ψ* are variance, spatial aspect ratio, wavelength of the sinusoidal component, and the phase offset of the sinusoidal component of the Gabor filter, respectively. We used both cosine and sine components of the Gabor filter. In our study we only varied the orientation; therefore, for simplicity of exposition, we only explicitly notate θ_i_ for parametrizing the Gabor filter in the following.

##### 2.2.4.2 Rectification

The output of the convolution contains both positive and negative values. Passing such positive and negative values through pooling can result in an attenuated response. Since we have both cosine and sine components of our Gabor filters, we deployed the energy model proposed for early visual cortex complex cells (Heeger, 1991; Carandini et al., 2005) according to

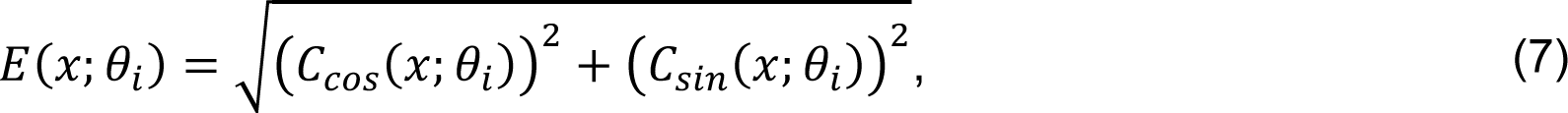

where *C_cos_* and *C_sin_* are the cosine and sine components of the Gabor filtered images, respectively.

##### 2.2.4.3 Normalization

We also included normalization with two goals in mind: First, to remove the sensitivity to local brightness; second, to prevent the rectification process from generating unbounded responses.

As divisive normalization is considered a canonical process across different cortical areas (Carandini & Heeger, 2012), we used a divisive form of normalization in our network according to

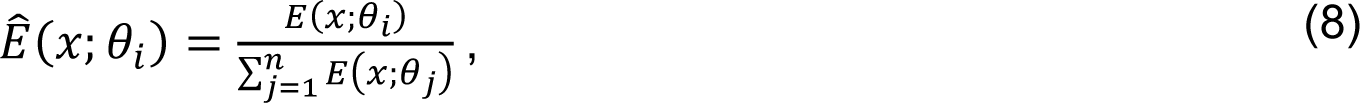

where j ranges across the n filter orientations.

##### 2.2.4.4 Spatial pooling

Spatial pooling is added to the network to aggregate normalized responses. This process provides a certain level of shift-invariance by abstracting the exact location of the responses in the pooling region. Similar mechanisms are observed in cortical areas (Ben et al. 2011). In our network, spatial pooling has been implemented using a low pass filter (i.e., binomial) and down sampling with factor two at each layer according to

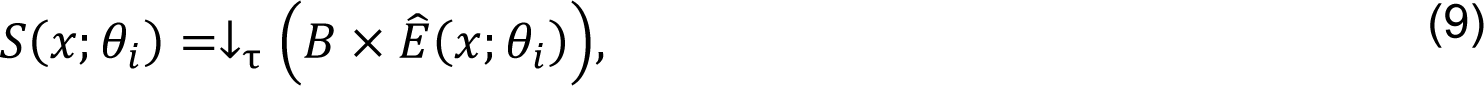

where *B* is a binomial lowpass filter and ↓_τ_ is spatial down-sampling. This spatial pooling is done at the end of each layer.

##### 2.2.4.5 Feature pooling

Finally, after the second layer, we included a feature pooling process. Our logic was that we created an abstraction for different features using repeated filtering (e.g., different orientations). Using the feature pooling process, we aim to create an abstraction for combined features (i.e., line-intersections). Another motivation for including feature pooling was to prevent the network from exploding, as after each filtering, the number of features will be doubled. We implemented the feature pooling process using a weighted summation of all the feature maps to yield

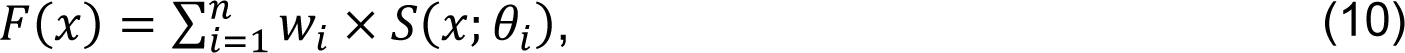

where *n* represents the number of features (e.g., for a network with 4 orientations in each layer and 2 layers *n* will be 16). The weights *W_i_* are trained to detect the intersection of the two lines and the target. Note that in our network, this feature pooling is only employed at the end of the second layer; however, for a network with more layers this mechanism may be exploited at the end of each layer (for further details see Hadji & Wildes, 2017). We complete processing at this layer by transforming the representation of the images into a vector format by sequentially concatenating the image rows (flattening). This flattened feature is fed to our MLP alongside initial gaze position data. Such flattening is typical when convolutional layer results are fed to fully connected layers and does not impact the signal content, even as it makes the format more amenable to fully connected processing.

#### 2.2.5 Fully connected layers

We used a physiologically inspired fully connected feed forward MLP to implement the visuomotor transformations for reaching toward the visual target. Figure 2C shows the schematic of the network architecture. The input to the MLP is comprised of two main types: 1) output of the CNN network that consists of extracted features from the input images, which has been flattened to facilitate further processing and 2) extra-retinal signals, which can consist of several different signals such as eye position, head configuration, etc., but in the current implementation is restricted to eye position These two types simply are concatenated for subsequent processing. The second layer is comprised of units that receive inputs from all the previous layer’s units and their activity is calculated as:

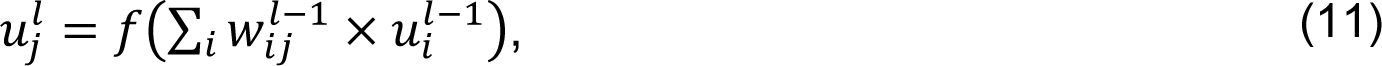

where, *u^l^_j_* is the activity of unit *j* at the current layer *l, W^l−1^_ij_* are learned connection weights from the *i^th^* to the *j^th^* unit, *u_i_^l−1^_i_* is the activity of unit *i* at the previous layer, and *f(x)* is the unit’s transfer function. In the current implementation, *l* = 2. Here, we considered the sigmoid transfer function to mimic the nonlinear transfer function of real neurons (Naka & Rushton, 1966) according to

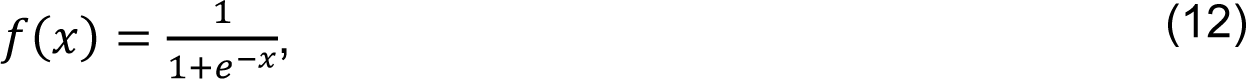

The units in the third layer provide the population coding of the reaching movement. As described in Section 2.2.3, these units are constrained to have cosine tuning. To impose such tuning behaviour, we constrained the activity of the third layer’s units by fixing the connection weights to the final read-out layer according to equations (2)-(4). The final layer provides 2D effector displacement read-out of the population coded Euclidean distance. We first trained a separate feed-forward network with one hidden layer to transform the population codes into 2D read-outs. Then, we fixed the connection weights between the third and final layer. Units of the final layer are purely linear.

### 2.3 Network training and testing

To provide realistic simulations and compare our model outputs to real data, we simulated the task employed in several previous experimental publications (Li et al. 2017; Bharmauria et al. 2020, 2021). As explained in Section 2.1, the task consists of 4 possible landmark locations and 8 possible landmark shifts. We extracted these landmark positions and their associated shifts as well as the target locations from the dataset. These data were used to create the encoding and decoding images. Additionally, in this experiment, 3D eye movements (horizontal, vertical and torsional components of orientation of the eye relative to space) were recorded (Bharmauria et al. 2020, 2021). For our simulations, we used the horizontal and vertical components to create the 2D initial and final gaze locations.

We made use of two types of datasets to train our network. First, a dataset was generated based on the available neurophysiological data (Bharmauria et al., 2020). We used approximately 25,000 trials for each of two monkeys, and we trained the network separately for each monkey. For each data point we had an encoding image (200×200), a decoding image (200×200), 2D initial eye position, and final gaze position to calculate the displacement of the eye. As described in the task, Section 2.1, an encoding image contains the initial eye fixation, target, and landmarks. Based on the task (Bharmauria et al., 2020), initial eye positions were distributed within 7°-12° from the centre of the screen, while the target and landmark locations were determined based on the neuron’s response fields. Target locations were distributed approximately within a rectangular range of 30-80° across both horizontal and vertical dimension. Exact target locations were determined based on the size and shape of a neuron’s response field (i.e., 4×4 or 7×7 grid and 5-10° apart). Landmark locations were selected from one of the oblique locations 11° apart from the target. The decoding images contained the visual landmark: in the exact same location as the encoding image or shifted 8° from the initial location. The direction of the shift was randomly selected from the 8 possible locations evenly distributed on a circle (for a visualization, see Figure 1).

To probe the network in more controlled, idealized conditions, we also generated simulated datasets for the combination of allocentric and egocentric information with 80,000 data points. We considered three scenarios: no allocentric (0% allocentric; final gaze location landed on the target location), purely allocentric (100% allocentric; final gaze location landed on the shifted target location), and a combination of allocentric and egocentric (30% allocentric; final gaze location landed between the target and shifted target location). Like the neurophysiological data, for each simulated data point we had an encoding image (200×200), a decoding image (200×200), 2D initial eye position, and final gaze position to calculate the displacement of the eye. For encoding images, we generated target and landmark locations randomly. These locations were uniformly distributed between (−40,40) and (−50,50) for landmark and target, respectively. For decoding images, landmark shifts were randomly chosen and varied in the range (−10,10). Similarly, initial gaze positions were generated randomly in the range (−10,10). All the values are generated in screen coordinates with (0,0) being the center of the screen.

We considered the Mean Square Error (MSE) between a monkeys’ final gaze position and the network output as the network loss for training,

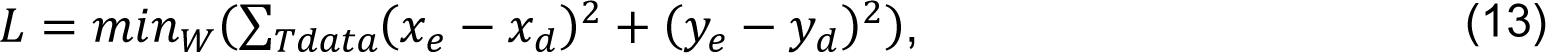

where *W = {W_1_, …, W_N_}* are the learned parameters, *Tdata* is training data, (*x_e_, y_e_*) is network’s estimated gaze location, and (*x_d_, y_d_*) is ground truth gaze location. The numerical value for *N* is specified below for each layer.

The values for the predetermined parameters in the first two layers of the CNN were as follows: we used four Gabor filters (7×7) with uniformly distributed orientations: *θ* ∈ 0°, 45°, 90°, 135°. All the other parameters were the same for all the Gabor filters (i.e., σ = 1, λ = 0.5, γ = 2, and ψ = 0). We used a down sampling of factor 2 for our spatial pooling layers. We used a sigmoid transfer function for the first layer of the feature pooling stage in the CNN (and the two first fully connected layers in our MLP). After the feature pooling, we flattened the images, which resulted in a vector with 4136×1 dimension. We used 44 units to code initial gaze location and 100 units for our hidden layer. For our motor population codes, we used 4 (for behavioural analysis) and 100 (for neural analysis) units. Overall, the learned parameters were 16 for the CNN and 535,040 and 13,376,000 for the MLP with 4 and 100 motor population units, respectively.

Training used the ADAM optimizer (Kingma & Ba 2015) with learning rate of 0.001 and batch size of 32. We divided each dataset into three sub-datasets: The first two subsets with proportion of 90 and 5 were used for training and cross-validation of the network. This separation was done to prevent overfitting. The final 5 precents were used for assessing the performance of the network against behavioural data.. We used R^2^ to quantify the explained variance in the training dataset by our network and evaluate the network performance. We stopped the training based on two criteria: 1) if the number of epochs reached 50, or 2) if the RMSE in the validation dataset stopped dropping significantly (i.e., gradients < 0.001).

### 2.4 Hidden unit analysis

#### 2.4.1 Fitting units’ response fields against spatial models

A key step in understanding how the brain implements the required coordinate transformation is determining the intrinsic coordinate frame of individual neurons. To be consistent with the published neural literature (Bharmauria et al. 2020, 2021), we used the same analytic method, i.e., fitting different spatial models to a neuron’s response field (Keith et al. 2009). The spatial model that yields the best fit (lowest residuals between the data and model) is taken as representative of the intrinsic coordinate frame of the neuron. Further description is provided in the supplementary method section.

In our task, and based on the neurophysiological findings, we considered three main reference points (Figure 4A): Target, final gaze, and shifted target locations. We also considered two possible coordinate frames, space vs. eye frames. To discriminate between these coordinate frames, we fit each unit’s response field against different spatial models (here six: Target in eye (Te), Final gaze in eye (fGe), Shifted target in eye (T’e), Target in space (Ts), Final gaze in space (FGs), and Shifted target in space (T’s)) using a non-parametric fit with a Gaussian kernel:

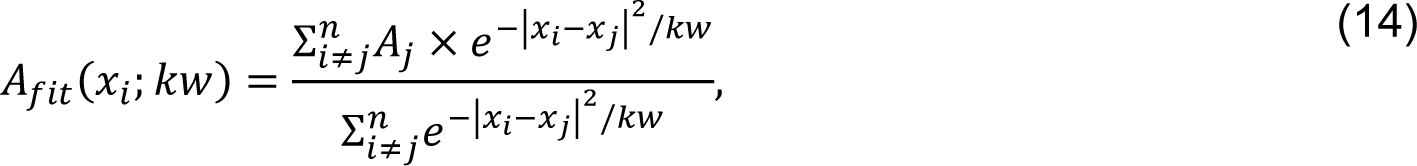

where *A_fit_* represents the fit prediction for a neuron’s activity, *A_i_* represents recorded activity for neuron *i, x* represents the position, and *kw* represents the Gaussian kernel bandwidth.

**Figure 4.**
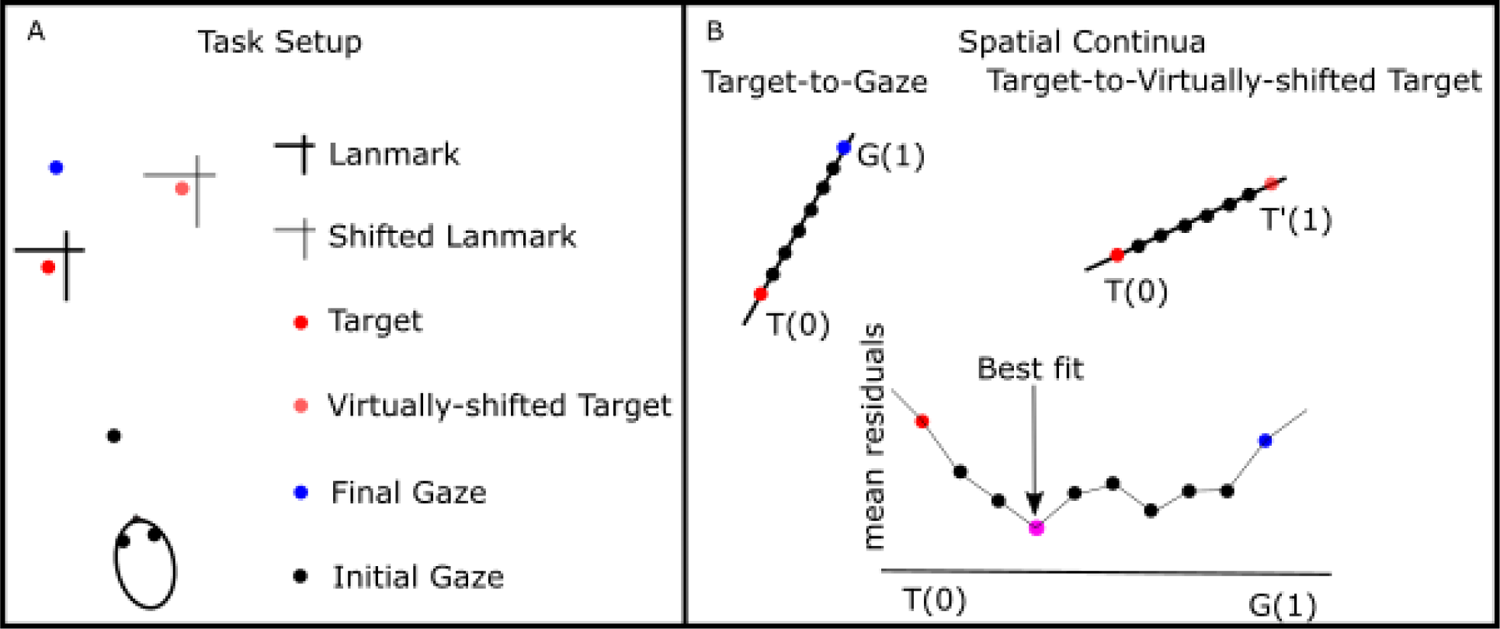
Spatial model fitting logic and procedure. A) Example of the task configuration: The box represents the real task under consideration, including both encoding (landmark and target) as well decoding (shifted landmark) images. B) Spatial continua: We selected three reference points to create spatial continua: Target, Gaze, and virtually shifted target locations. Using these three reference points we created two continua: Target-to-Gaze and Target-to-Virtually shifted Target. We performed model fitting across both continua and selected the model with the lowest mean residuals as the best model explaining intrinsic coordinate frames of the unit.

To quantify the goodness of the fit, Predictive Error Sum of Squares (PRESS; Keith et al., 2009) statistics were calculated. To do so, we calculated the fit for each trial based on the activity of the other trials. Then, we calculated error between the fit activity and the measure activity for each trial and calculated the mean squared difference for a given spatial model and kernel bandwidth. The model and kernel bandwidth that yielded the lowest residual is considered as the best fit. This kernel bandwidth is used for further analysis. Full details of this method are available elsewhere (Keith et al., 2009).

#### 2.4.2 Intermediate spatial models

Our previous results (Sadeh et al. 2015; Sajad et al. 2015, 2016; Bharmauria et al. 2020) suggested that canonical models are not always the best candidate to describe the neural response field, but instead intermediate models between canonical models should be considered. Additionally, based on our previous observations (Bharmauria et al. 2020, 2021), we designed our motor population layer to code information in eye coordinates. Consequently, in the following we only focus on the spatial models in eye coordinates and will not use subscripting. Figure 4A shows a schematic of the deployed task and Figure 4B demonstrates the two relevant continua. Therefore, we created two continua (Figure 4B): Target-to-gaze (T-G), and Target-to-virtually shifted target (T-T’). We considered 30 equally distributed steps for each continuum (with ten steps between the main reference points and 10 steps above each of the reference points). Furthermore, the rightmost panel in Figure 4B shows the residuals for the 10 steps between the target and gaze continuum. The model that yields the lowest PRESS residual is selected as the best model fit.

#### 2.4.3 Test for spatial tuning

The method of Section 2.4.2 can be used only if a unit’s activity is spatially tuned, i.e., selective for a particular set of target positions. Therefore, we tested for spatial tuning of each unit’s activity and excluded the spatially untuned units from our analysis. To test for spatial tuning, we shuffled the average firing rate data over the position data obtained from the best-fitting model. Then, we statistically compared the mean PRESS residuals distribution of the 200 randomly generated response fields with the mean PRESS residual distribution of the best fit model. A neuron’s activity is considered spatially tuned if the best fit PRESS residual fell outside of the 95% confidence interval of the distribution of the randomly shuffled mean PRESS.

## 3 Results

We evaluated our network in three ways. First, we evaluated general network performance by comparing the distribution of errors in its gaze output to errors in the training set, where the input was the same for both systems (i.e., same visual displays and initial eye positions). This comparison was done both for actual monkey data (Bharmauria et al. 2020) and simulated data, where we systematically manipulated allocentric vs. egocentric contributions to the data, with and without simulated noise. Second, we compared the allocentric-egocentric weighting produced by the network in each of these conditions to see if it would replicate the weighting of the training set (in particular, that of the actual monkey data). Third, we performed a detailed comparison between the activation of various units in the later ‘motor’ stages of our network against physiological data obtained from the frontal eye fields of the monkey in the same task (Bharmauria et al. 2020).

### 3.1 General performance: variable gaze errors, task dependence and noise

#### 3.1.1 Comparison between simulated and behavioral error distributions

The goal of this test was to ascertain if the network replicates the specific patterns and distributions of variable errors observed in monkeys on the same task. In the behavioural task, monkeys were rewarded if they performed a saccade within 8°-12° distance of the target (Bharmauria et al. 2020). Here, we used the data of the two monkeys to train and evaluate network performance. In all cases we tested the model on a portion of data that the network never saw during the training and validation phases. Note that for these tests we compared actual recorded gaze positions with gaze positions decoded from the MLP output layer of our model, trained on a different dataset of gaze end points derived from the same animal(s).

Figure 5A shows the final gaze locations (monkey #1 behaviour and our network’s prediction) for two randomly selected target locations (blue and red stars). Blue and red markers represent gaze locations for the first and second target respectively, while darker and lighter colors distinguish the monkey gaze from our network prediction with similar symbols indicating gaze positions for each trial. It is seen that, gaze end points are scattered around the target location (with higher noise for network’s prediction) but landed within the 8° or 12° radius, i.e., within the animal’s reward window. Figure 5B-C summarizes this observation across all trials in the same animal: Figure 5B shows the distribution of gaze error amplitudes (calculated as the angular distance between the end gaze point and target position) and Figure 5C shows the network generated data trained on a different set of data from the same animal. Supplementary Figure 1 shows a similar figure for the 2nd monkey. As can be seen, monkey data were very noisy, but the majority of the errors fell bellow 8°-12° of the visual target. Qualitatively, the trained network errors follow a similar distribution, except errors dropped off less precipitously outside the monkey’s reward window.

**Figure 5.**
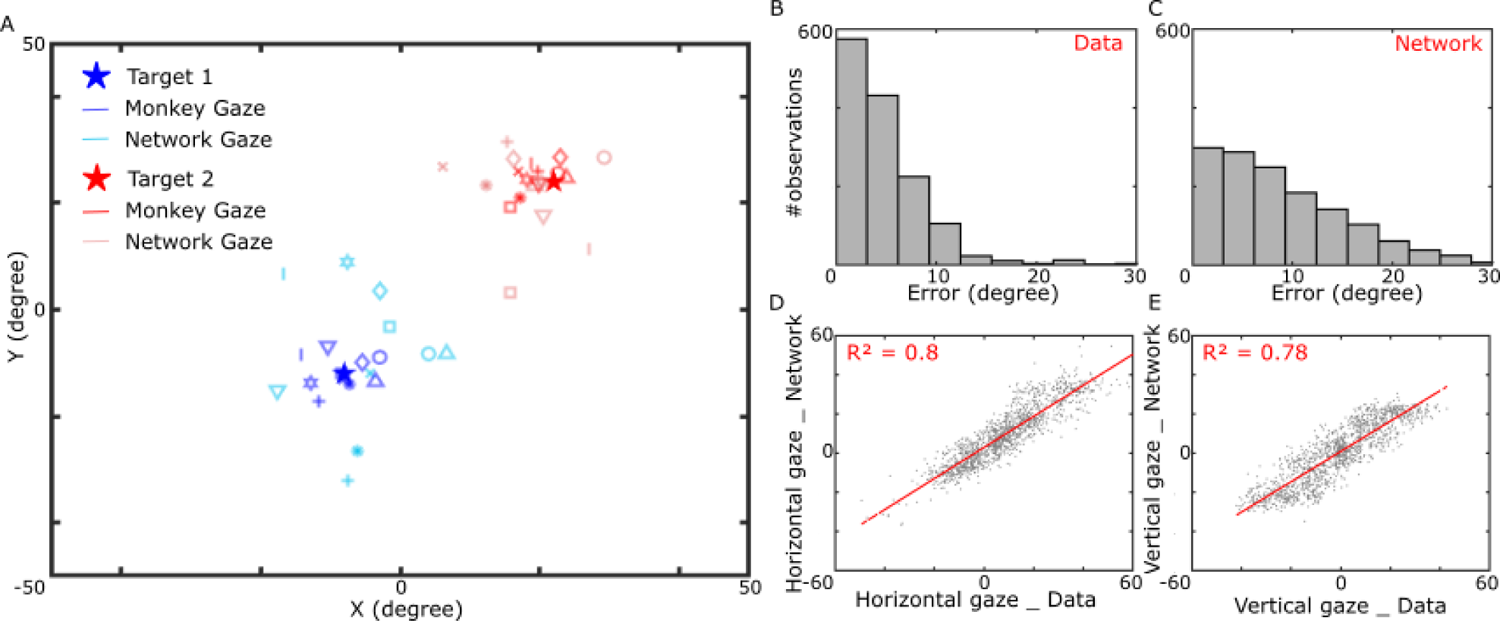
Network performance. A) Final gaze endpoints for two randomly selected target positions. Red and Blue stars show the target locations. Darker colors represent monkey’s data while lighter color represent our network’s prediction. Points with the same marker represent the same trial (simulated vs. recorded monkey data). Our network followed the monkey behaviour for both target locations. However, our network’s predictions are more scattered (higher variability) compared to the monkey data. B) Distribution of gaze endpoint errors observed from the monkey data. The errors are mainly below 8-12°. C) Distribution of gaze endpoint errors generated by our network. The majority of errors are below 8-12° similar to the data. D-E) Regression analysis of our network gaze endpoints and the observed data. Our network explained approximately 80% of the data’s variability for both horizontal and vertical directions.

To quantify these observations, we calculated the goodness of fit of the network output to the actual data using R^2^, which assesses the ability of our network to explain the data’s variability (Figure 5 D-E) Broken down into components, the model explained 80% of the horizontal variance and 78% of the vertical variance in this animal (refer to supplementary Figure 1 for similar result of the second animal). Overall, our network explained 80% of the data’s variability in the first animal, and 75% in the second animal (Table 1, rows 1-2). In the following sub-section, we assess how well the model generalizes to other task situations, and the possible role of noise in these data fits.

**Table 1.**
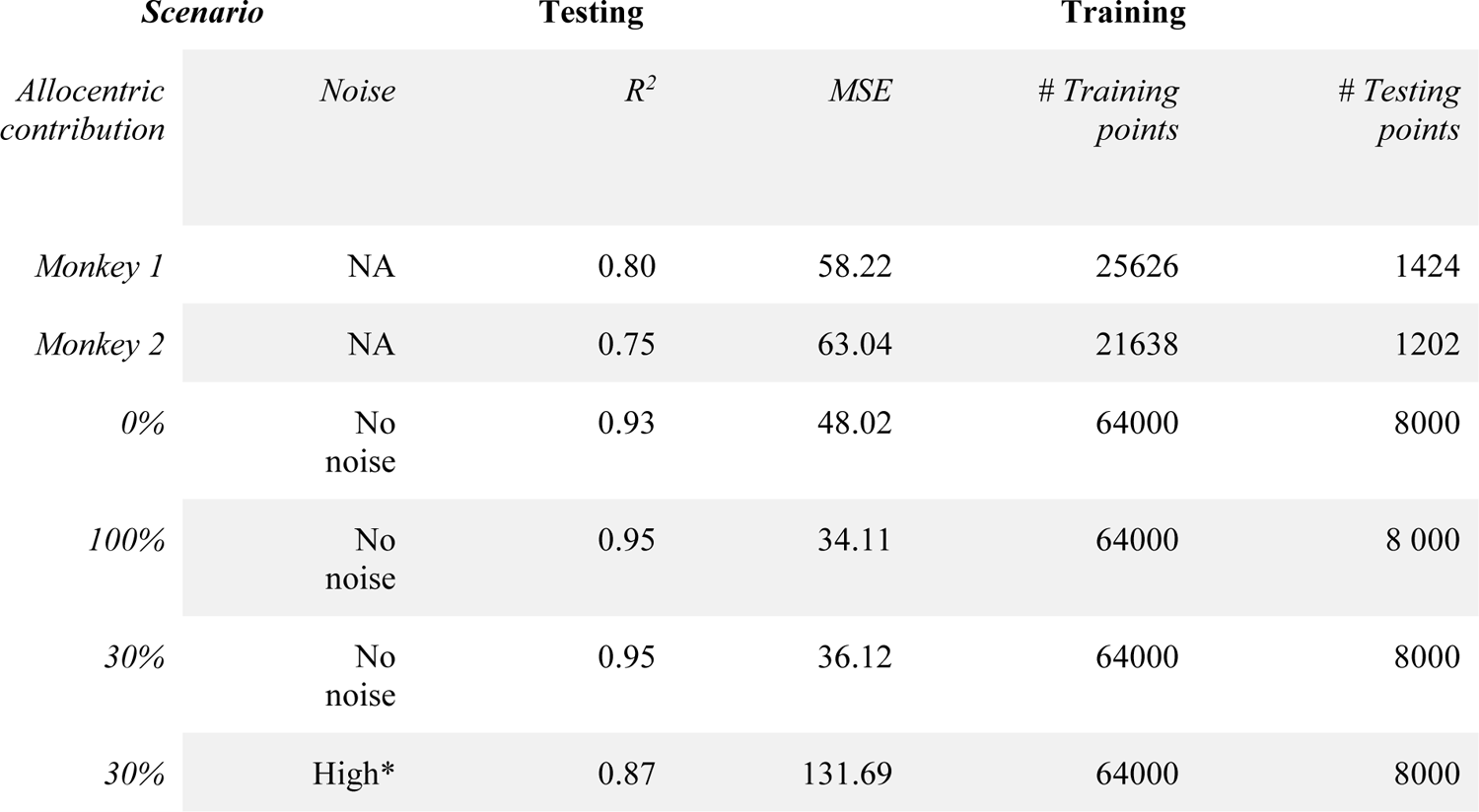
Evaluation of our network performance under different scenarios and comparison to monkey data.

#### 3.1.2 Generalization and Noise

The purpose of this analysis was to evaluate 1) if our network outputs could generalize to different task scenarios, and 2) the influence of noise in the training set. For the first goal, we created 3 different synthetic training sets, with different levels of allocentric-egocentric weighting: 100% - 0%, 0%-100%, and 30%-70% (similar to that observed in experimental studies). In other words, the final gaze positions were located on the target, on the virtual shifted target, or 30% shifted toward the virtual shifted target respectively, in the training set. No noise was present in these datasets. Table 1 (rows 3-5) summarize the network’s performance for these scenarios, providing R^2^ values for ideal vs. actual performance (Note that for this test, different datapoints were used than those used in the training set). It is seen that all three task situations, the network is capable of predicting a considerable amount of the data’s variability (> 90%).

The performance of the model was better in these idealized datasets compared to the performance of the model trained on actual data. We hypothesized that this was very likely because there was noise in the monkey data unrelated to the task. To test this hypothesis, we created a fourth synthetic training set: again 30%-70% allocentric-egocentric, but with added noise (random gaze variations) similar to that seen in the data. This manipulation decreased the explained variability from 95% to 87% (Table 1, row 6), supporting the notion that much of the unexplained variance in our data-trained model was due to input noise rather than some failure of the model to simulate systematic behavior. The latter is examined in the next section.

### 3.2 Systematic Performance: Allocentric-Egocentric Weighting

The key feature of allocentric-egocentric integration in behavioral studies is that when these cues conflict, humans and monkeys perform as if weighing between them. The weighting of allocentric cues tested so far has ranged from 30-50%, depending on the experimental conditions (Neggers et al. 2005; Byrne & Crawford 2010; Fiehler et al. 2014; Klinghammer et al. 2017; Li et al. 2017; Lu & Fiehler 2020). This result presumably reflects an optimization process, where usually egocentric and allocentric cues would tend to agree with each other, but with different levels of reliability and noise (Byrne et al. 2007; Körding et al. 2007; Byrne & Crawford 2010; Lew & Vul 2015; Klinghammer et al. 2017; Aagten-Murphy & Bays 2019). Here, we directly tested if our network learned to perform such integration.

#### 3.2.1 Network replicates the influence of a landmark shift in primate gaze behaviour

We now specifically assess whether the gaze endpoints produced by our trained networks replicate the influence of the landmark shift when actual monkey data are used in the training set. The influence of the shift is quantified by the component of the gaze endpoints (d) along the axis between the target (T) and direction of the landmark shift (T’) (Bharmauria et al. 2020). As this parameter quantifies the contribution of allocentric information in reaching, it is taken as the allocentric weight (AW): AW = 0 means that no allocentric information is used (i.e., the gaze endpoints were in close vicinity of the memorized target), while AW = 1 means that gaze is completely influenced by allocentric information (i.e., gaze endpoint shifted with the same amplitude and direction as landmark shifts). To directly compare our network’s prediction with monkey’s behaviour, we trained the network on each animal’s dataset and then calculated AW for both the training datasets (Figure 6 A-B) and the network’s output (Figure 6 C-D). Notably, in all cases (both training sets, and both networks) the gaze endpoints are shifted toward the shifted landmark with average AW ≈ 0.33. We observed higher variance in the network output, which we postulate is mainly related to noise in the training sets. We explore this matter further in the next section.

**Figure 6.**
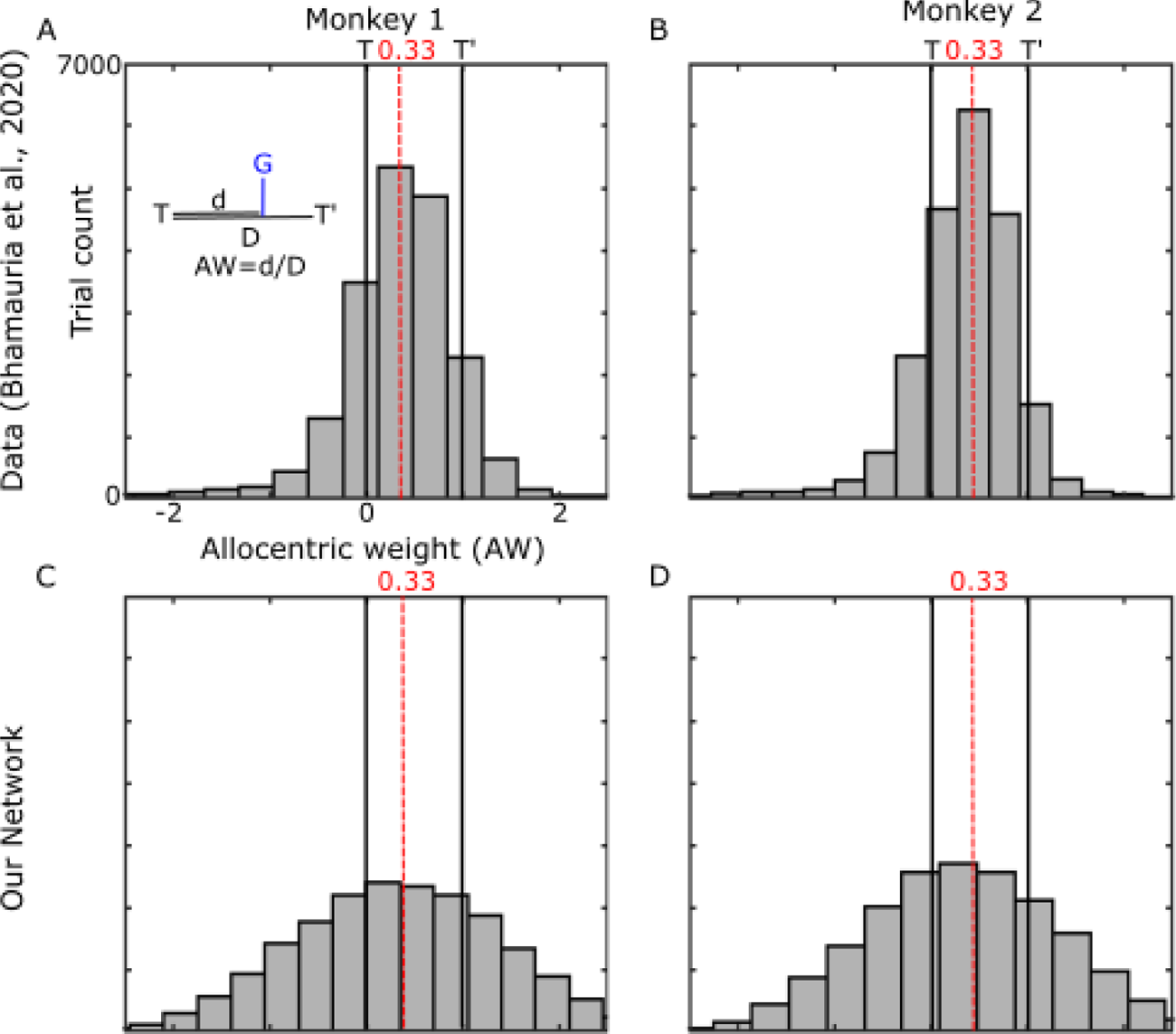
Allocentric weight based on Monkey’s behaviour. A-B) Allocentric weights for two monkeys. Shifting the landmark caused a scatter of gaze endpoint along the T-T’ axis. However, on average gaze endpoints were shifted 33% toward the shifted landmark location. C-D) Allocentric weights calculated based on our network predictions. Both networks show similar behaviour as the data: 33% shifted gaze endpoints in the direction of the shifted landmark.

#### 3.2.2 Allocentric-Egocentric Integration in Simulated Training sets

To explore how our model generalizes to other training sets and reconsider the role of data noise in this process, we performed a similar analysis (Figure 7) on models trained on the synthetic datasets described above, with 0%, 100%, 30% allocentric weighting, and 30% plus noise.

**Figure 7.**
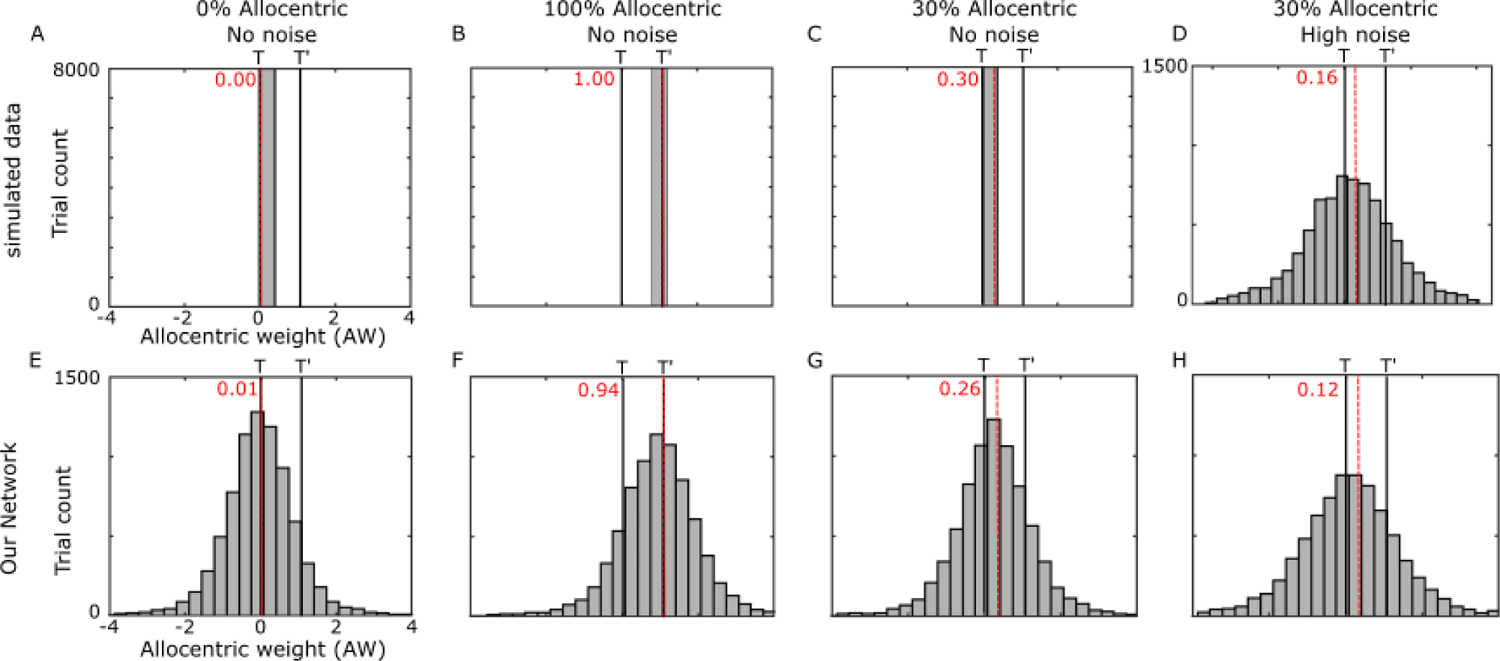
Allocentric weight based on simulated dataset. A-D) Allocentric weights for different simulated datasets. A) No allocentric contribution with no noise: the final gaze location landed on the remembered target location, B) full allocentric contribution with no noise: the final gaze location landed on the virtual target location, C) partial allocentric contribution (30%) with no noise: final gaze location landed between the remembered target and virtual target location. The percentage determines the amount of shift toward the virtual target. D) partial allocentric contribution (30%) with added noise (Gaussian): final gaze location distribution has a Gaussian form, with mean between the remembered target and virtual target location. The percentage determines the shift of the mean toward virtual target location. E-H) Our network’s replication of the final gaze locations for different simulated datasets where the differences arise from different selected experiment parameters (i.e., allocentric weight, and noise). In all scenarios, the network was able to adapt the location of the final gaze location based on the desired allocentric contribution (0%, 100%, and 30%). As expected, the network’s predicted gaze locations were noisy for all scenarios with the highest noise when the training dataset was noisy.

Figure 7 illustrates that our network learned different integration of allocentric and egocentric information dependent on the training dataset. For instance, varying the contribution of allocentric information from 0% (Figure 7A) to 100% (Figure 7B) resulted in a shift of the gaze endpoint distribution from being centered on 0 (Figure 7E) to being centered on 1 (Figure 7F). Similarly, including noise in the training dataset (Figure 7C vs. Figure 7D) resulted in higher variance in the network’s generated gaze endpoints (Figure 7G vs. Figure 7H). Based on these simulations, we conclude that 1) the model is able to achieve arbitrary levels of allocentric-egocentric integration, depending on the input and 2) again, it is likely that noise in the training set causes differences from the monkey data, rather than some fundamental limitation in the model.

### 3.3 Allocentric-Egocentric coordinate frames: MLP output unit analysis

We aimed to produce a model that can be compared directly to observed neurophysiological data. Specifically, we tested if the model was able to recreate a similar distribution of spatial coding observed in actual FEF data in the same task (e.g., Sajad et al. 2015; Bharmauria et al. 2020). Here, the key observations are that 1) FEF motor responses code gaze in an eye-centred frame of reference, 2) this code was partially shifted toward the shifted landmark, and 3) the influence of the landmark shift was fully integrated into the eye-centered motor responses of saccade.

To assess if our network shows similar results, we created a simulation of the neurophysiology experiments. To provide a uniform dataset of target/landmark combinations we generated a simulated training / testing dataset that contained a uniform distribution of target locations across the encoding-decoding images. This latter design was essential to ensure that we have enough units responsive to different target locations. (Note that our actual behavioral data missed many of these points, but networks trained on the behavioral data gave similar results to those described below.) To generate the final gaze positions, we created a Gaussian distribution for egocentric (Target) and allocentric (Virtually-shifted target) information. We then created a distribution for the final gaze position using a weighted summation of allocentric and egocentric information (Bayesian integration; Byrne et al. 2007; Körding et al. 2007; Klinghammer et al. 2017). For our integration, we considered 33% weights for the allocentric information. This weight selection resulted in a similar distribution to the observed data (as shown in the previous section). We also added noise to replicate the monkey data (similar to row 6 in Table 1). Finally, we sampled from this distribution to generate the gaze position for each trial. We used this dataset to train our network. Then, we created a dataset where target location incrementally changes across the images to assess the intrinsic coordinate frames of hidden units in our motor population layer. We defined the motor population layer to resemble the FEF code (Baharmauria et al., 2020, 2021), as explained in the methods. For these simulations, we simulated 100 MLP output neurons with 2D directional tuning distributed evenly across 360°.

#### 3.3.1 Response Fields and Intrinsic coordinate frames of MLP output units

Recall that we constrained our output to represent FEF motor neurons. To this aim, we implicitly forced the motor population units to have cosine tunings with open ended response fields. Here, we first confirm that our motor units behave as they are designed. Figure 8A shows an example motor response field, from a unit in motor population layer, in the absence of a landmark shift. The plotting conventions are the same as those used for real neural data in our previous papers, i.e., circle size indicates the neural ‘firing rate’ for a randomly selected subset of the individual trials, and the color code represents the non-parametric fit made to the full dataset. Both the individual trial locations and fit are plotted in the coordinate system that provided the best overall fit for this unit: future gaze relative to initial eye position (see methods and below for further explanation). Similar to our actual FEF motor response recordings (Sajad et al. 2015; Bharmauria et al. 2021) we observed that only a subset of our neurons are spatially tuned (similar to Bharmauria et al., 2021) and that spatially tuned units showed ‘open ended response field (as expected).

**Figure 8.**
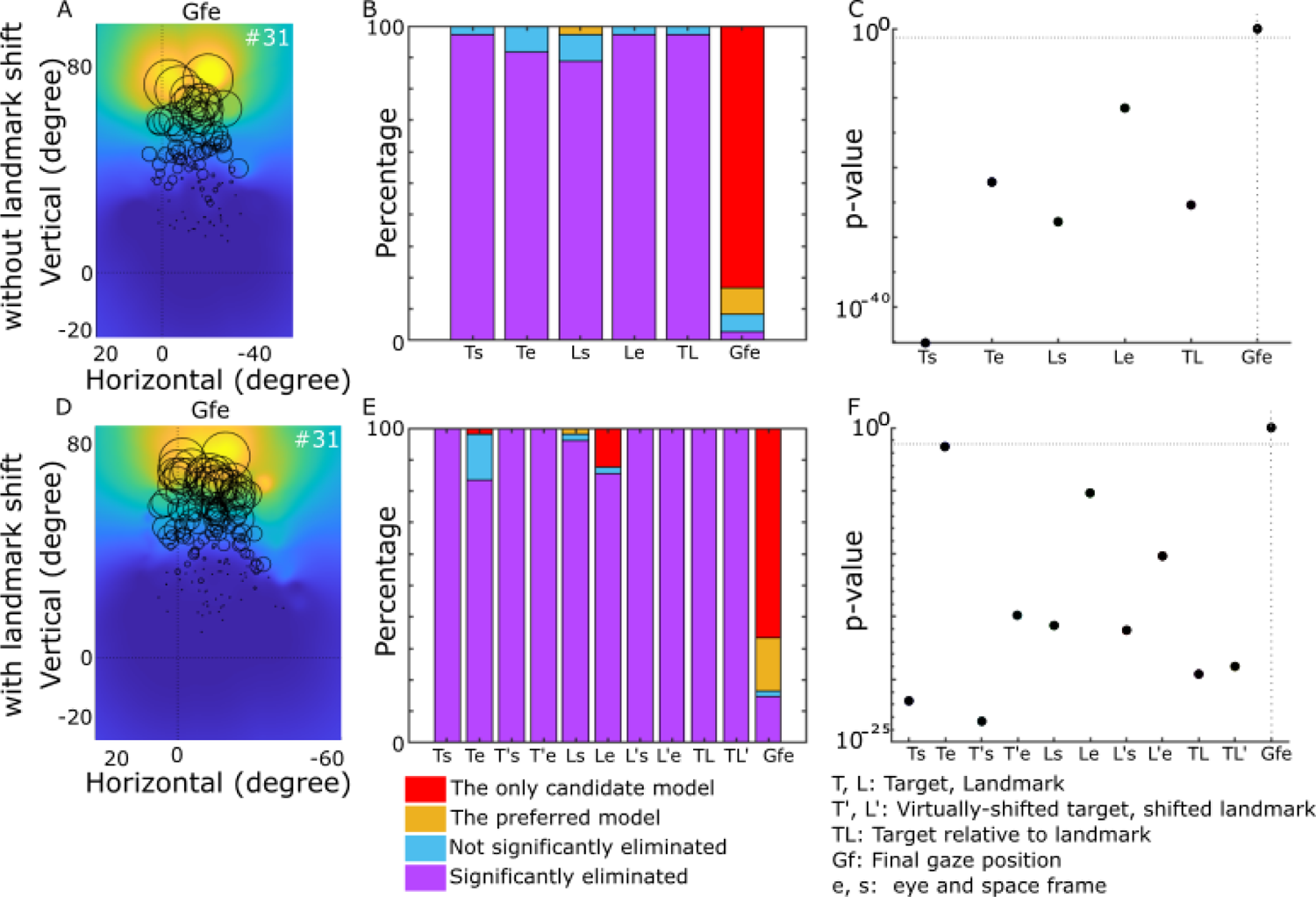
Intrinsic coordinate frame analysis for motor population’s units. A—C) without landmark shift: A) the response field of a motor unit. This unit has an open-ended response field with and upward preference. Cooler colors represent lower activity and warmer colors represent higher activity. Our non-parametric fit revealed that this unit significantly (p < 0.05) prefers a gaze-in-eye coordinate frame. The prediction of our fit is presented by black circles where the diameter indicates the activity predicted by our fit. We reduced the number of circles (randomly) for illustration purposes. B) Summary of the preferred coordinate frames for all 34 spatially tuned motor neurons. The tested models were Target in space (Ts), Target in eye (Te), Landmark in space (Ls), Landmark in eye (Le), Target in landmark (TL), and Gaze in future eye (Gfe). The majority of neurons (∼83%) significantly (p < 0.05) preferred gaze-in-eye coordinate frames. C) population analysis of units coordinates frames. The population analyses were performed by a t-test of mean residuals for each model relative to the best model fit. When coordinate frames were tested at the population level, the entire population significantly (p < 0.05) preferred gaze-in-eye coordinates. D-F) Similar analysis as A-C but in the presence of the landmark shift. Here we had 48 spatially tuned neurons. In addition to the previously mentioned spatial models, models related to landmark shifts were tested: Virtually-shifted target in space (T’s), Virtually shifted target in eye (T’e), Shifted landmark in space (L’s), Shifted landmark in eye (T’e), and Target in shifted landmark (TL’).

The next step is to investigate the coordinate frames our units deploy to code information. To test the intrinsic coordinates of our simulated neural population, we used the same methods used in our previous physiological studies (Bharmauria et al. 2020, 2021). In brief, we calculated the residuals between the individual data points and fits made in each coordinate system. Only units that showed significant spatial tuning were selected for further analysis (n = 34 in the absence of landmark shift). We tested Target-in-space (TS), Target-in-eye (Te), Landmark-in-space (Ls), Landmark-in-eye (Le), Target-relative-to-landmark (TL) and future Gaze relative to initial eye position (Gfe). We found that the majority of individual units (∼83%) showed a significant preference for Gfe coordinates, and no neurons showed a significant preference for the other models (Figure 8B). Similarly, the entire population showed a significant (P < 0.05) preference for gaze-in-eye coordinates (Figure 8C), similar to Sajad et al. (2015), where there was no landmark.

We repeated the same analysis in the presence of the landmark shift (Figure 8 D-F), where 48 of the 100 output units showed significant spatial tuning. Here, additional models were included to account for the shifted landmark position (L’) and the virtual position of the target relative to the landmark (T’L’). However, these results showed a similar pattern: response fields were very similar (Figure 8D), a few units showed a significant preference for Le but the majority (∼67%) showed a significant preference for Gfe (Figure 8E) and the entire population significantly preferring future gaze-in-eye coordinates (Figure 8F). In short, these results replicated those recorded from the monkey FEF in the presence of a landmark shift: preservation of the basic eye-centered gaze code (Bharmauria et al. 2020). But as in the latter study, this comparison of ‘cardinal’ models was not sensitive enough to detect the influence of the landmark shift. For that, we turned to a more sensitive analysis based on intermediate coordinate frames, as shown below.

#### 3.3.2 Influence of landmark shift on MLP output unit response fields

In monkeys, FEF motor responses are modulated by a shift in the landmark, specifically causing a shift toward landmark-centred coding without altering the basic response field or gaze code (Bharmauria et al. 2020). To see if a landmark shift produced a similar influence in our simulated data, we first plotted unit response fields for different landmark shifts. A typical example is shown in Figure 9. Neural activity is represented by the heat map, with an asterisk (*) placed at the peak of the response field, plotted in Gfe coordinates. and indicate the coordinates of landmark shifts on top of each box (in parentheses). The middle panel in figure 9 shows the response field for no shift condition (0,0), and the other panels are arranged congruently with the landmark shift direction (i.e., up for up-shift etc.). Again, the unit activity resembles an open-ended response field. At first glance, the Gfe response field appears to be very similar for each landmark shift, consistent with the notion that this unit is primarily coding gaze. However, the peak of the response field appears to shift with the landmark in some directions (e.g., for left and right shifts in this case) but not other shift directions. This result was typical, but other units showed direction-dependencies (of the cue shift) on both the magnitude and direction of their response field shifts (see supplementary Figure 2). To understand the overall landmark influence on this population, we re-examined the underlying coordinates of these response fields, using a more sensitive test (Bharmauria et al. 2020, 2021).

**Figure 9.**
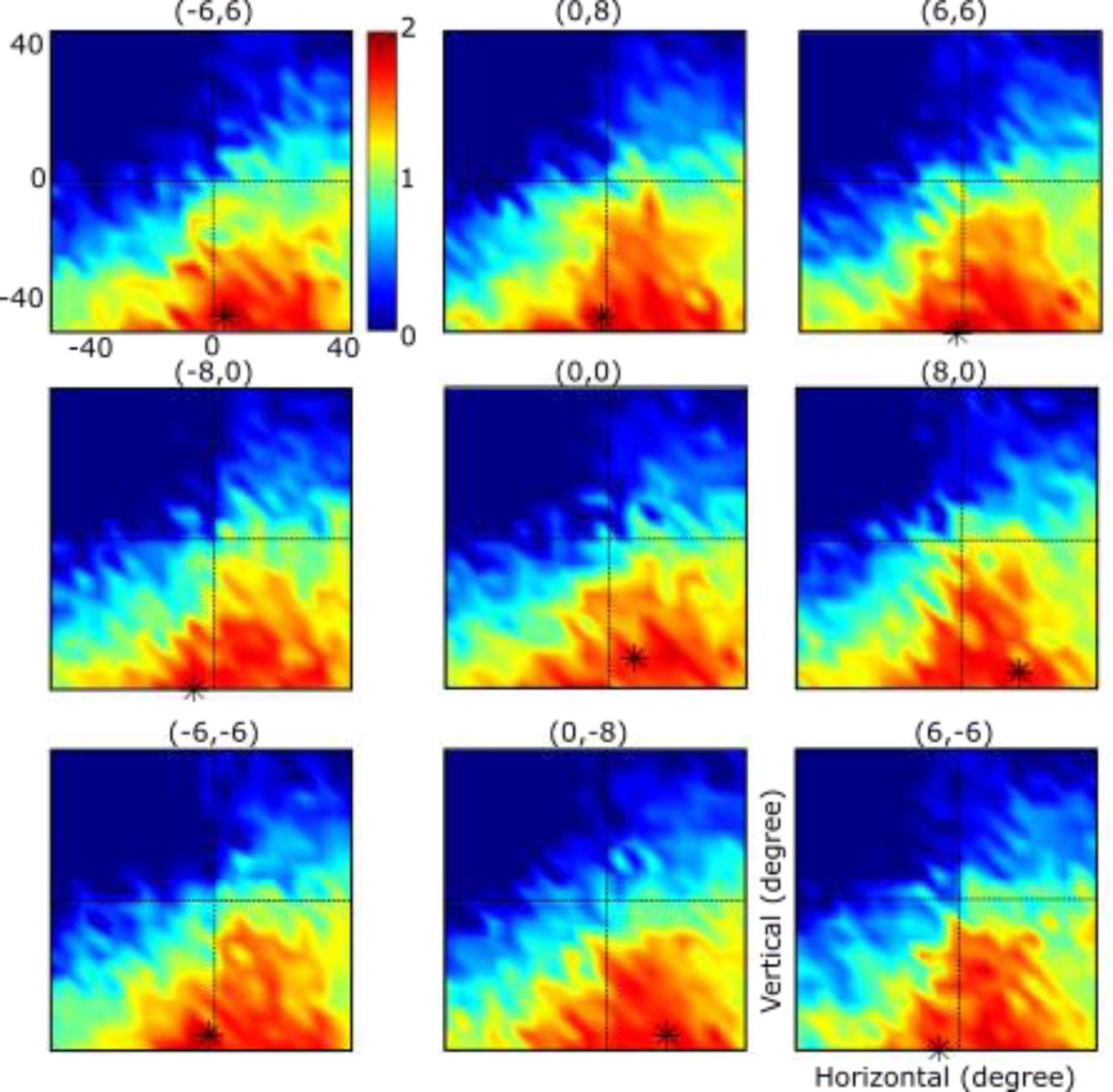
A unit’s response field for different landmark shifts. Each panel shows the response field of the same unit for different landmark shifts. The coordinates of landmark shifts are provided on top of each box (in parenthesis). Response fields are plotted in gaze coordinates. The peak of activity is indicated by an asterisk (*) in each box. Following the asterisk location, varying the landmark shift resulted in a correspondingly shifted response field in some directions indicating the possibility of shifted intrinsic coordinate frames.

#### 3.3.3 Influence of landmark shift on intrinsic coordinate frames of MLP output units

In the next step, we performed a similar ‘intermediate frame’ analysis as used in our previous neurophysiology studies (e.g., Bharmauria et al., 2020). Specifically, we used non-parametric fits (as described in Section 2.4) to detect the best fits for each spatially tuned unit along two spatial continua: Target to shifted target (T-T’) and Target to Gaze (T-G). T-T’ provides a continuous measure of the influence of the landmark shift on the target representation, where T’ would be a virtual target fixed to the shifted landmark (Bharmauria et al., 2020). T-G measures the degree to which each unit encodes variable gaze errors (Sajad et al. 2015). Each continuum was divided into 10 steps between the two ‘cardinal’ models with 10 more steps beyond each. The point yielding the lowest fit residuals represents the best intrinsic coordinate frame.

Figure 10 provides a direct comparison of actual FEF motor responses (top row) vs. our network’s motor population layer (bottom row). The first two columns of Figure 10 A-D show fits along T-T’ continuum. The first column shows an example FEF neuron that codes information in an intermediate coordinate frame that is shifted 3 steps in the direction of T’ (the landmark shift), with a very similar simulated unit (also shifted 3 steps) shown below. The second column shows corresponding frequency histograms for the physiological and simulated data. The distribution of the physiological fits was broader compared to the simulated data fits but they both show a shift to the right, i.e., in the direction T’. For the physiological data the shift had a mean of 29% (median 20%) toward T’. The simulated data, 3 steps (30%) were shifted slightly less but in the same direction (mean = 20%, median = 12%, p =3.3e-7, Wilcoxon signed ranked test). These shifts, along the T-T’ continuum, are qualitatively consistent with the observed 33% allocentric shift in the behavioural data (see Section 4.2.1). These results suggest that, like the actual data, our MLP output unit coordinates shifted partially with the landmark toward coding T’.

**Figure 10.**
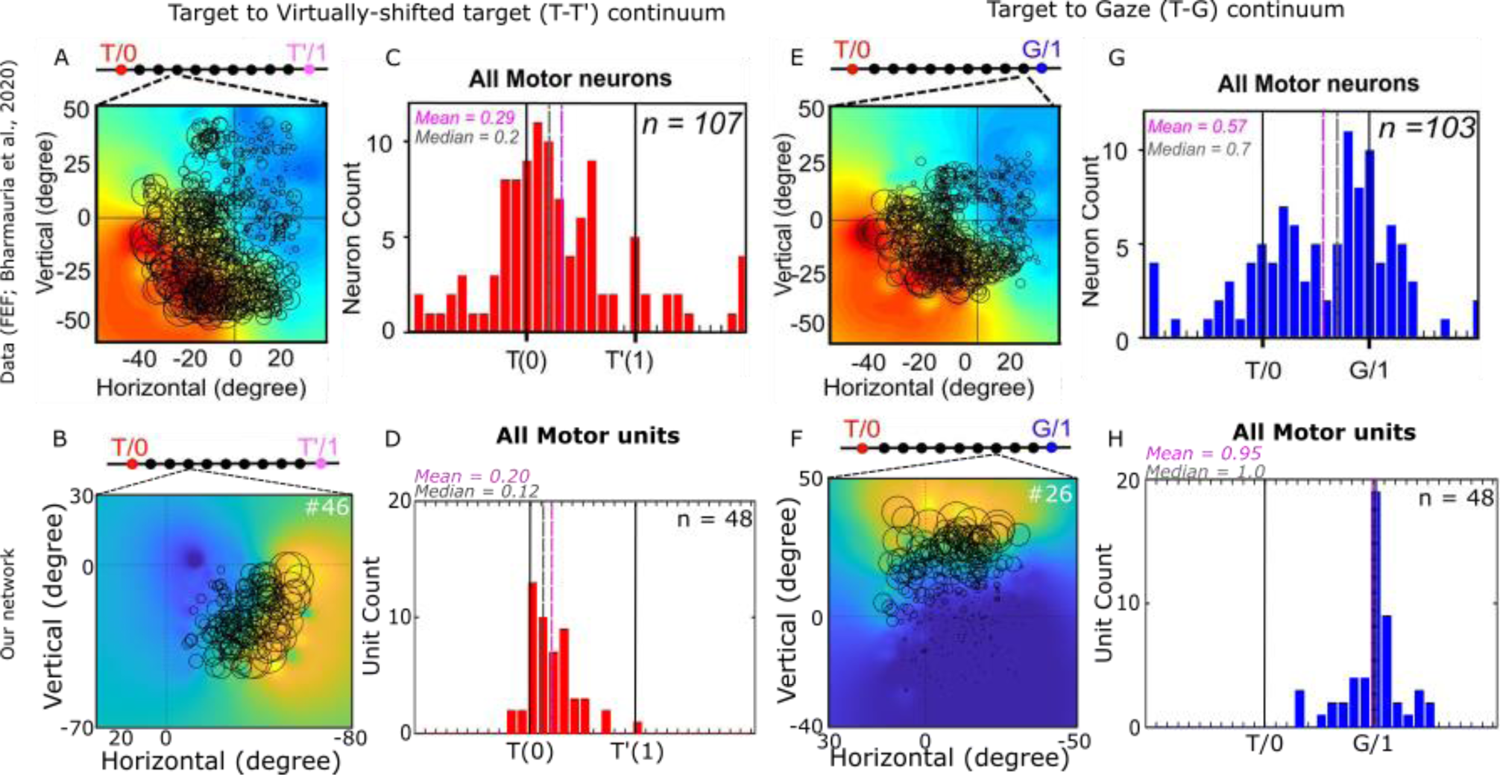
Motor neuron’s coordinate frames. Model hidden units at the motor population layer showed similar coding mechanisms as the neurophysiology data. A-D) Model fits in allocentric coordinates (i.e., along T-T’ continuum). A-B) Fit model for a sample unit in FEF [adapted from Bharmauria et al., 2020 with permission] (A) and in our motor population layer (B). In both example neurons, the intrinsic coordinate frame was 3 steps away from the target toward the shifted target. C-D) Distribution of the intrinsic coordinate frames when fit along T-T’ continuum. Similar to FEF neurons (C), motor population coding in our network (D) was biased toward target coordinates when fit along the Target-to-Shifted target coordinates with a partial shift toward the shifted target (30%). E-H) Model fits in egocentric coordinates (i.e., along T-G continuum). E) Fit model for an example neuron. In this neuron the best model is 1 step before the final gaze model. F) Fit model for a unit in our motor population layer. G) Distribution of the intrinsic coordinate frames of motor neurons in area FEF (Bharmauria et al., 2020). The majority of motor neurons coded information in Gaze coordinates. H) Distribution of the intrinsic coordinate frames of units in our motor population layer. Similar to FEF data, the majority of motor neurons coded information in Gaze coordinates. [E and G were adapted from Bharmauria et al., 2020 with permission]

The last two columns of Figure 10 E-H provide a similar analysis for the T-G continuum. The example response fields (3^rd^ column) were shifted 90% and 70% from T toward G in the real and simulated data respectively. The fourth column shows the distribution of best fits for the real and simulated data. In this case, the real data showed a bimodal distribution with a smaller peak near T and a larger peak near G, and an overall mean and median of 57% and 70% respectively. The simulated data (lower panel) showed a significant shift (p =2.6e-10, Wilcoxon signed ranked test) toward G that was larger on average with a mean of 0.95% and median of 100%. However, this dataset appears to correspond to only the ‘G’ peak in the physiological data. Accounting for this, the dominant peaks of both datasets indicated near pure gaze coding. Overall, these results indicate that the model, like most actual FEF responses, show a coordinate shift in the direction of the landmark while continuing to code gaze relative to initial eye orientation. Although this might sound contradictory, in the next section we show how these two results can be reconciled.

#### 3.3.4 Integrated Landmark Influence in the Final Motor Response

In the previous section, we showed that similar to FEF motor neurons, our motor population units code information in both egocentric (T-G) and allocentric coordinates (T-T’). An important question is whether these two codes are integrated or independent. To answer this question, we correlated our allocentric and egocentric codes. As illustrated in Figure 11A, these codes could be completely independent (vertical or horizontal lines), multiplexed but uncorrelated (shifted lines), or correlated (diagonal lines). In the physiological data recorded previously (Bharmauria et al. 2020), these measures were uncorrelated in memory responses (not shown), but became significantly correlated in the motor response, suggesting an integrated motor code (Figure 11B). We tested if the same was true in our simulated data. Although the simulated data had a smaller distribution, we observed a significant allocentric-egocentric correlation (Figure 11C; slope = 0.2370 ± 0.17, R^2^ = 0.2731, p = 0.050) in our motor output layer. This result suggests that our network replicates the observed behaviour in monkeys FEF motor responses, i.e., an allocentric influence that is integrated into the egocentric motor response to directly influence gaze behavior.

**Figure 11.**
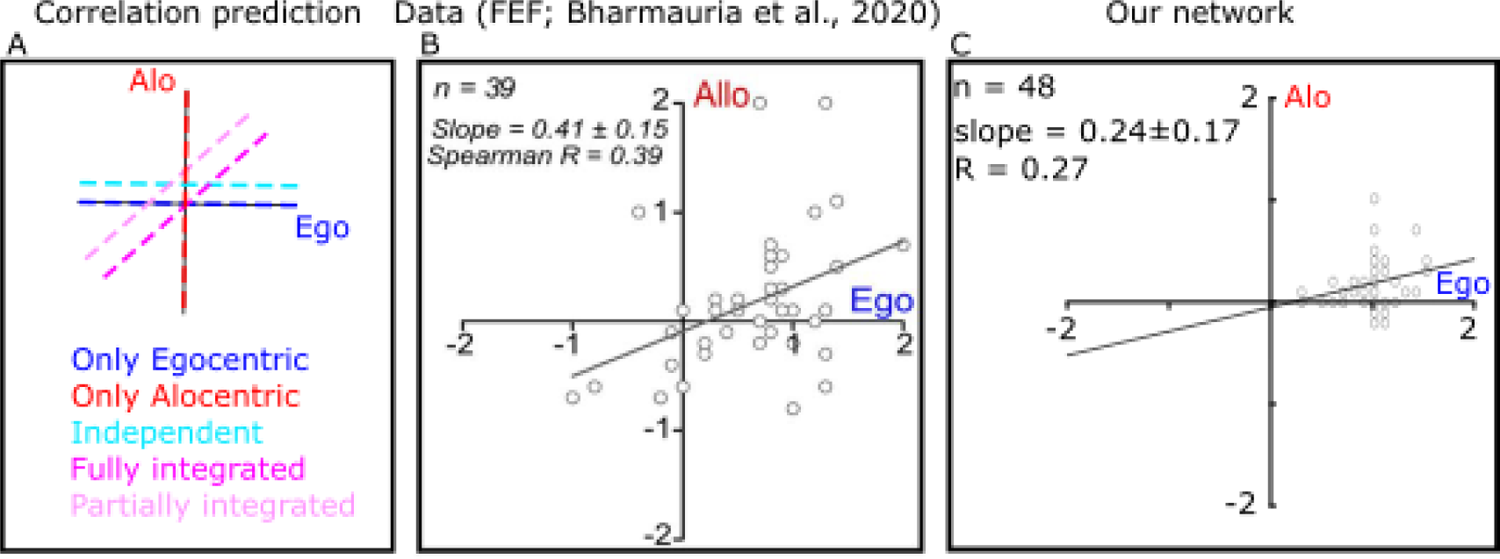
Integration of allocentric and egocentric information. A) Correlation prediction for different allocentric and egocentric integration scenarios. B) Allocentric information was integrated into FEF motor codes. When allocentric coding is examined against egocentric coding, it showed significant correlation [adapted from Bharmauria et al., 2020 with permission]. C) Similar to the FEF data (Bharmauria et al., 2020), our motor population layer showed integrated allocentric information into the motor codes (significant correlation between allocentric and egocentric coding; slope = 0.2370 ± 0.17, R = 0.2731, p = 0.050).

## 4 Discussion

In this study, we developed and validated a novel neural network framework of allocentric-egocentric integration for goal directed movements. The model used physiologically constrained inputs and outputs and was validated through comparison of 1) network output with known monkey gaze behavior in cue-conflict task and 2) its hidden unit’s activity with FEF neural activity recorded in this same task (Baharmauria et al., 2020). To implement a general theoretical framework, we modelled the visual system as an analytically specified CNN and the sensorimotor transformation for gaze as a MLP. The network (i.e., MLP weights) was trained on synthetic and actual data from a task where a landmark shifted relative to the saccade target. We observed that 1) our network generates saccade vectors comparable to observed monkey gaze as well as simulated datasets with different allocentric-egocentric contribution, with and without noise, 2) the network replicates the allocentric-egocentric weighting in different datasets, and 3) our motor units code gaze in an eye-centred frame; this code was partially shifted toward the shifted landmark and the influence of the landmark shift was fully integrated into the eye-centered motor responses of saccade, similar to FEF motor neuron responses. To our knowledge, this is the first network model that combines complex visual system properties with a sensorimotor transformation to replicate observed behavioural and neural data.

### 4.1 Neural Network Models in Sensorimotor Neuroscience

Neural network approaches have long been deployed to understand the underlying mechanisms for sensorimotor transformations (for reviews, see Pouget & Snyder 2000; Battaglia-Mayer et al. 2003; Blohm et al. 2009). The general approach is to train an analogous network using similar input-output as observed in sensorimotor tasks, evaluate network output and performance against experimental data (as we have done here) and then analyze hidden unit properties to understand how the brain might do this. Early studies modeling sensorimotor transformation in 2D found that varying initial eye, head, and hand positions results in gain modulation of unit’s activity (Zipser & Andersen 1988; Salinas & Abbott 1995; Xing & Andersen 2000). Analysis of hidden layers revealed that hidden units encode information in purely gaze-centered coordinate frames, while the information is coded in intermediate coordinate frames or as shifting response fields in motor output layers. Extended from 2D modeling, a 3D neural network investigation revealed new properties generalizing the previous findings (Smith and Crawford 2005; Blohm et al. 2009). The authors observed fixed input-output relationships within each unit and each layer. These fixed input-output relationships acted as local coordinate transformation modules. The global transformation at the network level was implemented by combining these local transformations, weighted in a gain-field like fashion (Blohm et al. 2009). These new properties emerged as a result of the nonlinearities inherited form the 3D coordinate transformations. Other studies have extended such models to include recurrent connections, ‘memory’ and spatial updating of the remembered visual goal during eye movement (Keith et al., 2010).

The above-mentioned models played a crucial role in understanding the available data as well as providing prediction for further experiments. However, these studies have two shortcomings. First, in all the previous studies, the complexity of visual system is ignored: The visual stimulus is oversimplified into a ‘dot’ that is represented as a hill of activity in a retinal topographic map. This oversimplification results in the inability of such networks to model complex stimuli (e.g., crossing lines, or objects). Thus, previous models are not capable of explaining the presentation of allocentric information and their role in sensorimotor transformations. Second, the majority of the previous studies treated sensorimotor transformations as a feedforward network that resulted in a lack of temporal dynamics. Here, we attempted a first step toward addressing the first of these two limitations (visual complexity), as discussed in the following sections.

### 4.2 Current Approach

To create and validate a model that would be useful for neurophysiologists, we followed principles established in previous studies described above (Zipser & Andersen, 1998; Blohm et al., 2009; Keith et al., 2010). In particular, we constrained the inputs and outputs of our network to resemble the known physiologically of the modeled system. Since our aim was to reconstruct the early visuomotor transformations from visual cortex to early two-dimensional (2D) eye centered motor codes in frontal cortex (Bharmauria et al. 2000, 2001) we simplified the inputs as 2D eye-centred images and 2D eye position signals based on the eye-position coding observed in S1 (Wang et al., 2007). Likewise, we modeled the output layer to represent the open-ended response fields observed in the FEF (Bruce & Goldberg, 1985; Sommer & Wurtz, 2006; Sajad et al. 2015, 2016). This particular input-output configuration allowed us to both train and compare our model outputs to a known dataset (Bharmauria et al. 2000, 2001).

In contrast to previous visuomotor networks (which processed a single ‘dot’ stimulus), our model had to encode a 2D visual image containing multiple features and then somehow represent / integrate egocentric and allocentric information derived from this image. To do this, it was necessary to combine the complexity of the visual system (Schenk 2006; Thaler and Goodale, 2011; Geirhos et al., 2017; Rajalingham et al., 2018) with the with feed-forward properties of a sensorimotor transformation (Smith & Crawford, 2005; Blohm et al., 2009; Crawford et al., 2011). For this purpose, we found it useful to separate the network into a representational stage (i.e., the visual system) and a sensorimotor transformation stage (Crawford et al. 2011).

To model the visual system, we used a CNN. But unlike previous models (Geirhos et al., 2017; Rajalingham et al., 2018; Schrimpf et al., 2018; Kar et al., 2019; Lindsay 2021) we didn’t train our CNN but instead used an analytically constrained architecture to be consistent with the known physiology. Specifically, we built our CNN based on the concept of repeated filtering: “filter-rectify-filter”. Our network consisted of two identically designed layers in which each layer consists of convolution, rectification, normalization, and pooling. To replicate the properties of cortical simple and complex cell, we deployed 2D oriented Gabor filters for our convolutions (Carandini et al., 2005). Following the biological evidence suggesting that learning occurs at higher cortical levels (Serre et al., 2005), we introduced a trainable feature pooling layer to construct the final output from CNN with the required attributes (here landmark and visual target). This design enabled us to construct a highly interpretable model of the visual system.

To model the sensorimotor system, we used the outputs of our CNN as inputs for a fully connected feedforward MLP. In line with sensorimotor transformations studies, this network was left fully trainable. As noted above, previous studies have shown that such feedforward networks are able to produce striking similarities with observed neural activities in areas involved in sensorimotor transformations (Zipser & Andersen, 1998; Blohm et al., 2009; Keith et al., 2010). For this purpose, it is advantageous that our network is highly interpretable, i.e., it is not a black box as is often the case for many artificial neural networks.

Finally, we trained and validated our model against real data. We trained our network using both idealized datasets and behavioral data from a recent neurophysiology experiment (Bharmauria et al, 2020). For training purposes, we decoded behavior (saccade vectors) from the motor output, based on physiological realism and previous network studies (Smith & Crawford, 2005; Blohm et al., 2009). We chose to model saccades because of their simplicity and the availability of relevant data, but our model can be generalized to other visuomotor behaviors by adapting its input-output structure, for example to resemble the reach system (Blohm et al. 2009).

### 4.3 Optimal Allocentric-Egocentric Integration in Behavior

In an early study that combined experimental data with statistical modeling, Byrne and Crawford (2010) showed that reach movements were biased toward shifted landmark when participants were instructed to reach to memorized visual targets in the presence of a visual landmark (presented as vibrating dots on the corners of an invisible square). The amount of shift was influenced by the actual relative reliability of gaze versus landmark positions, as well as the perceived stability of the visual landmarks: higher vibration resulted in less reliance on the landmark information. More recent experiments with more realistic scenes, replicated the basic result (i.e., aiming shifted with landmarks) but showed that the magnitude landmark influence was determined by different factors such as the number of shifted objects (Fiehler et al., 2014; Klinghammer et al., 2015) and scene consistency (Klinghammer et al., 2017). Similar effects have been observed in monkey gaze behaviour (Li et al., 2017; Bharmauria et al., 2020).

Here, we showed that our network is capable of reproducing the observed monkey saccades with good agreement (R2≈75%-80%; table 1, rows 1 and 2), as well as various other scenarios with different allocentric-egocentric weightings (Figures 6, 7). Our network reproduced similar allocentric weights as reported for monkey’s data (Figure 6), but with wider distribution (Figure 7). Some of the differences with real data might be accounted for by the existence of noise in sensorimotor systems (Körding & Wolpert 2006; Alikhanian et al., 2015; Abedi Khoozani & Blohm, 2018). Consistent with this, increasing the noise reduced the ability of the network to explain data variability (table 1, rows 3-6). This perhaps agrees with a recent report that the addition of noise to deep neural nets improves correspondence with actual response fields in early visual cortex (Jang et al. 2021).

How does the brain implement allocentric-egocentric integration? At a computational level, the predominant view in the field is that the brain uses probabilistic inference to estimate the reliability of ambiguous sensory signals for optimal integration (Ernst & Banks, 2002; Alais & Burr, 2004; Pitcow & Angelaki, 2014). In this view, the brain estimates each signal contribution for the optimal integration (e.g., for allocentric-egocentric weighting) based on the statistical properties of sensory signals (where higher signal variability results in lower contribution). This can be derived online through probabilistic neural codes (Ma et al., 2006) or based on learned contextual cues (Mikula et al. 2018). For example, Byrne and Crawford (2010) explained their reach data using a maximum likelihood estimator based on the actual reliability of egocentric versus allocentric cues combined with a landmark stability heuristic based on prior experience. Here, our neural network model learned to extract such rules from the training set, consistent with the suggestions that error-based learning rules naturally implement probabilistic inference (Orhan & Ma, 2017). But to understand how this is done, it is necessary to look within actual neural populations (Pitcow & Angelaki, 2014).

### 4.4 Simulation of Neurophysiological Results

As noted above, we coded the motor output of our network using the same motor population coding mechanisms as seen in the FEF (Bruce & Goldberg, 1985; Sommer & Wurtz, 2006; Sajad et al. 2015, 2016). This was choice was made to 1) impose physiological realism within the model and 2) allow direct comparisons of our network model with actual FEF data. To test if our motor output units generate similar properties as their counterpart motor neuron in FEF, we performed three analyses similar to neurophysiology studies. First, we confirmed that our motor population units show open ended receptive fields and code information in a gaze-centered coordinate frame (Figure 8). These observations are in line with previous observation of dominate gaze-centered coding in motor neurons (Sajad et al., 2015, 2016). While, the open receptive field arose due to our design, the gaze-centered coding emerged from the training process. Second, and more importantly, when we examined the intermediate coordinate frames, we observed that our units use a range of intermediate coordinate frames between target and gaze to code the information as reported for FEF and SEF motor neurons. Third, examining the influence of the landmark shift, we observed that shifting the landmark resulted in shifted receptive fields (Figure 9). In particular, we observed that the coordinate frames of our units were partially shifted toward the virtually shifted target (Figure 10) representing an allocentric coding in our network. When we evaluated the allocentric-egocentric information, we found a strikingly similar partial integration as reported in SEF and FEF. On the whole, these results suggest that we successfully created a model that can closely follow current neurophysiological reports (Bhamariua, 2020). This was a crucial goal for the current project. These observations suggest that the underlying mechanisms of the proposed network might have close similarities with brain mechanisms for allocentric-egocentric integration.

### 4.5 Potential Interpretability of the network processes

As suggested above, the striking similarities between our unit’s activity and their physiological counterparts suggest that our network might use similar mechanism as the brain to implement the required processes. This implies that interpretability of our network is essential. We argue that our network is highly interpretable by design. First, all the units of our CNN except, the feature pooling layer, are determined analytically; thus they are fully interpretable. For the example, the feature pooling is fully mathematically measurable: a product of all the feature responses in the final maps determining crossing points of the two lines as well as the target location. The next and most challenging task will be to uncover the mechanisms governing the transformation *between* the input-output in our MLP. In particular, what happens in the trainable layers between our analytically defined visual system layers and motor output layer. For this purpose, it should be possible to use both analytic solutions arising from computer science (e.g., Hadji & Wildes 2017; Monga et al. 2020; Zhao & Wildes 2021) and ‘neurophysiological’ techniques like those employed in the past (Zipser & Andersen, 1998; Smith & Crawford, 2005; Blohm et al., 2009). In this way, our visual-motor model can be used as a ‘quasi experimental model’, in parallel with studies on the real brain, for example to understand how egocentric and allocentric signals are integrated in goal-directed movements.

### 4.6 Limitations and Future Directions

Despite its success in replicating current neurophysiological and behavioural observation, this study is only the first step in modeling the integration of complex visual stimuli for sensorimotor transformations. To extend its usefulness beyond the current task (allocentric-egocentric cue conflict in the gaze system) several steps need to be taken. First, it should be possible to train this model on other 2D datasets (different visual stimulus configurations etc.) to see how well it generalizes and investigate other questions involving the use of complex visual stimuli for movement control. Second, it should be possible to extend this model to include 3D geometry of sensorimotor transformation (and including other extra retinal signals such as head position), multisensory (e.g., somatosensory) inputs of target and hand position. As emphasized by many studies, modeling the 3D linkage of eye-head-hand is crucial for a full understanding of sensorimotor transformations (for a review see Crawford et al., 2011). Third, another limitation of the current version of the model is that it does not include any temporal dynamics. Temporal dynamics are essential for for quantifying the progression of sensory coding through memory delay signals and into motor coding in such tasks (Bharmauria et al. 202, 2021). Including recurrent connection to the current model addresses this limitation. Finally, the current version of the model is appropriate for detecting simple stimuli such as 2D crosses or dots and therefore is not generalizable to datasets that include more realistic 3D objects. However, our studies in dynamic texture recognition show that incorporating additional CNN layers enables the current CNN model to recognize more complicated patterns (Hadji & Wildes 2017).

### 4.7 Conclusions

We implemented and evaluated a neural network model that provides both the capacity for encoding relatively complex visual features and sensorimotor transformation for goal-directed movements. We trained this model on real and synthetic datasets involving saccade generation in the presence of allocentric landmarks. We showed that our network replicates the reported behaviour and generate the observed neural activities in FEF areas. These results suggest that our framework provides an analytic toolbox to better understand the interaction of allocentric and egocentric information for goal-directed movements. We further propose that this network can be generalized to model other complex visuomotor tasks. Building such toolboxes is necessary to facilitate our further understanding of the underlying mechanisms for performing sensorimotor coordinate transformations where the stimulus is not simply a ‘dot. Since spatial transformations ubiquitously underly most brain processes, this toolbox has potential for application in fields as diverse as reaching to grasp, posture/balance control, visual navigation, decision making, as well as their analogs in computer vision and robotics.

## Acknowledgements

This project was supported by Vision: Science to Applications Program (to P.A.K); the Canada Research Chair Program (to J.D.C.); an NSERC Grant (to R. P. W.). We thank Sohrab Soleimani who did preliminary work with an earlier version of the model.

## References

1. Aagten-Murphy, D., Bays, P. M. (2019). Independent working memory resources for egocentric and allocentric spatial information. PLOS Comput Biol. 15: e1006563.

2. Abedi Khoozani, P., Blohm, G. (2018). Neck muscle spindle noise biases reaches in a multisensory integration task. J Neurophysiol 120: 893–909. doi:10.1152/jn.00643.2017.

3. Alais, D., & Burr, D. (2004). The Ventriloquist Effect Results from Near-Optimal Bimodal Integration. Current Biology, 14, 257–262. https://doi.org/10.1016/j.cub.2004.01.029

4. Alikhanian, H., de Carvalho, S. R., Blohm G. (2015). Quantifying effects of stochasticity in reference frame transformations on posterior distributions. Front Comput Neurosci 9: 82. doi:10.3389/fncom.2015.00082.

5. Andersen, R. A., & Buneo, C. A. (2002). Intentional Maps in Posterior Parietal Cortex. Annual Review of Neuroscience, 25(1), 189–220. https://doi.org/10.1146/annurev.neuro.25.112701.142922.

6. Ball, K., Smith, D., Ellison, A., & Schenk, T. (2009). Both egocentric and allocentric cues support spatial priming in visual search. Neuropsychologia, 47(6), 1585–1591. https://doi.org/10.1016/j.neuropsychologia.2008.11.017

7. Baker, C. L., & Mareschal, I. (2001). Processing of second-order stimuli in the visual cortex. Progress in Brain Research, 134, 171–191. https://doi.org/10.1016/S0079-6123(01)34013-X

8. Battaglia-Mayer, A., & Caminiti, R. (2018). Parieto-frontal networks for eye–hand coordination and movements. In Giuseppe Vallar, H. Branch Coslett (Ed.), Handbook of Clinical Neurology, 499–524. https://doi.org/10.1016/B978-0-444-63622-5.00026-7

9. Bharmauria, V., Sajad, A., Li, J., Yan, X., Wang, H., & Crawford, J. D. (2020). Integration of Eye-Centered and Landmark-Centered Codes in Frontal Eye Field Gaze Responses. Cerebral Cortex (New York, N.Y.: 1991), 30(9), 4995–5013. https://doi.org/10.1093/cercor/bhaa090

10. Bharmauria, V., Sajad, A., Yan, X., Wang, H., & Crawford, J. D. (2021). Spatiotemporal coding in the macaque supplementary eye fields: Landmark influence in the target-to-gaze transformation. ENeuro, 8(1), 1–23. https://doi.org/10.1523/ENEURO.0446-20.2020

11. Blohm, G., & Crawford, J. D. (2007). Computations for geometrically accurate visually guided reaching in 3-D space. Journal of Vision, 7(5), 4. https://doi.org/10.1167/7.5.4

12. Blohm, G., Keith, G. P., & Crawford, J. D. (2009). Decoding the cortical transformations for visually guided reaching in 3D space. Cerebral Cortex, 19(6), 1372–1393. https://doi.org/10.1093/cercor/bhn177

13. Byrne, P. A., & Crawford, J. D. (2010). Cue Reliability and a Landmark Stability Heuristic Determine Relative Weighting Between Egocentric and Allocentric Visual Information in Memory-Guided Reach. Journal of Neurophysiology, 103(6), 3054–3069. https://doi.org/10.1152/jn.01008.2009

14. Carandini, M, & Heeger, D. (2012). Normalization as a canonical neural computation. *Nature Reviews Neuroscience*, (November), 1–12. https://doi.org/10.1038/nrn3136

15. Carandini, M. (2006). What simple and complex cells compute. Journal of Physiology, 577(2), 463–466. https://doi.org/10.1113/jphysiol.2006.118976

16. Carandini, M., Demb, J. B., Mante, V., Tolhurst, D. J., Dan, Y., Olshausen, B. A., Gallant, L. G., & Rust, N. C. (2005). Do We Know What the Early Visual System Does?, 25(46), 10577–10597. https://doi.org/10.1523/JNEUROSCI.3726-05.2005

17. Caruso, V. C., Pages, D. S., Sommer, M. A., & Groh, J. M. (2018). Beyond the labeled line: Variation in visual reference frames from intraparietal cortex to frontal eye fields and the superior colliculus. Journal of Neurophysiology, 119(4), 1411–1421. https://doi.org/10.1152/jn.00584.2017

18. Chen, Y., Byrne, P., & Crawford, J. D. (2011). Time course of allocentric decay, egocentric decay, and allocentric-to-egocentric conversion in memory-guided reach. Neuropsychologia. https://doi.org/10.1016/j.neuropsychologia.2010.10.031

19. Chen, Y., Monaco, S., & Crawford, J. D. (2018). Neural substrates for allocentric-to-egocentric conversion of remembered reach targets in humans. European Journal of Neuroscience, 47(8), 901–917. https://doi.org/10.1111/ejn.13885

20. Crawford, J. D., Henriques, D. Y. P., & Medendorp, W. P. (2011). Three-Dimensional Transformations for Goal-Directed Action. Annual Review of Neuroscience, 34(1), 309–331. https://doi.org/10.1146/annurev-neuro-061010-113749

21. Danjo, T. (2020). Allocentric representations of space in the hippocampus. Neuroscience Research, 153, 1–7. https://doi.org/10.1016/j.neures.2019.06.002

22. Ernst, M. O., & Banks, M. S. (2002). Humans integrate visual and haptic information in a statistically optimal fashion. Nature, 415, 429–433. https://doi.org/10.1038/415429a

23. Fiehler, K., Wolf, C., Klinghammer, M., & Blohm, G. (2014). Integration of egocentric and allocentric information during memory-guided reaching to images of a natural environment. Frontiers in Human Neuroscience, 1–12. https://doi.org/10.3389/fnhum.2014.00636

24. Fukushima, K., Harada, C., Fukushima, J., & Suzuki, Y. (1990). Spatial properties of vertical eye movement-related neurons in the region of the interstitial nucleus of Cajal in awake cats. Experimental Brain Research, 79(1), 25–42. https://doi.org/10.1007/BF00228871

25. Gabor, D. (1946). Theory of communication. Journal of the Institution of Electrical Engineers, 93, 429–441.

26. Geirhos, R., Janssen, D. H. J., Schütt, H. H., Rauber, J., Bethge, M., & Wichmann, F. A. (2017). Comparing deep neural networks against humans: object recognition when the signal gets weaker. BioRxiv. http://arxiv.org/abs/1706.06969

27. Goodfellow, I., Bengio, Y., & Courville, A. (2016). Deep learning, The MIT Press.

28. Hadji, I., & Wildes, R. P. (2017). a spatiotemporal oriented energy network for dynamic texture recognition. In Proceedings of the IEEE International Conference on Computer Vision (ICCV).

29. Harandi, M., Sanderson, C., Shen, C., & Lovell, B. (2013). Dictionary learning and sparse coding on Grassmann manifolds: An extrinsic solution. In Proceedings of the IEEE International Conference on Computer Vision (ICCV).

30. Heeger, D. I. (1991). Nonlinear model of Neural responses in Cat Visual Cortex. In M. L. and J. Movshon (Ed.), Computational Models of visual Processing (pp. 119–134). MIT Press.

31. Hubel, D. H., & Wiesel, T. (1962). Receptive fields, binocular interaction and functional architecture in the cat’s visual cortex. The Journal of Physiology, 160, 106–154.

32. Hubel, D., & Wiesel, T. (1968). Receptive fields and functional architecture of monkey striate cortex. The Journal of Physiology, 215–243. https://doi.org/papers://47831562-1F78-4B52-B52E-78BF7F97A700/Paper/p352

33. Jang, H., McCormack, D., & Tong, F. (2021). Noise-trained deep neural networks effectively predict human vision and its neural responses to challenging images. PLOS Biology, 19(12), e3001418. https://doi.org/10.1371/journal.pbio.3001418

34. Kakei, S., Hoffman, D. S., & Strick, P. L. (2001). Direction of action is represented in the ventral premotor cortex. Nature Neuroscience, 4(10), 1020–1025. https://doi.org/10.1038/nn726

35. Kakei, S., Hoffman, D. S., & Strick, P. L. (2003). Sensorimotor transformations in cortical motor areas. Neuroscience Research, 46(1), 1–10. https://doi.org/10.1016/S0168-0102(03)00031-2

36. Kalaska, J. F., Scott, S. H., Cisek, P., & Sergio, L. E. (1997). Cortical control of reaching movements. Current Opinion in Neurobiology, 7(6), 849–859. https://doi.org/10.1016/S0959-4388(97)80146-8

37. Kar, K., Kubilius, J., Schmidt, K., Issa, E. B., & DiCarlo, J. J. (2019). Evidence that recurrent circuits are critical to the ventral stream’s execution of core object recognition behavior. Nature Neuroscience, 22(6), 974–983. https://doi.org/10.1038/s41593-019-0392-5

38. Keith, G. P., DeSouza, J. F. X., Yan, X., Wang, H., & Crawford, J. D. (2009). A method for mapping response fields and determining intrinsic reference frames of single-unit activity: Applied to 3D head-unrestrained gaze shifts. Journal of Neuroscience Methods, 180(1), 171–184. https://doi.org/10.1016/j.jneumeth.2009.03.004

39. Keith, G. P., Blohm, G., & Crawford, J. D. (2010). Influence of saccade efference copy on the spatiotemporal properties of remapping: A neural network study. Journal of Neurophysiology, 103(1), 117–139. https://doi.org/10.1152/jn.91191.2008

40. King, W. M., Fuchs, A. F., & Magnin, M. (1981). Vertical eye movement-related responses of neurons in midbrain near interstitial nucleus of Cajal. Journal of Neurophysiology, 46(3), 549–562. https://doi.org/10.1152/jn.1981.46.3.549

41. Kingma, D. P., & Ba, J. L. (2015). Adam: A method for stochastic optimization. 3rd International Conference on Learning Representations, ICLR 2015 - Conference Track Proceedings, 1–15.

42. Körding, K. P., & Wolpert, D. M. (2006) Bayesian decision theory in sensorimotor control. Trends Cogn Sci 10: 319–326. https://doi.org/10.1016/j.tics.2006.05.003

43. Klier, E. M., & Crawford, J. D. (1998). Human oculomotor system accounts for 3-D eye orientation in the visual-motor transformation for saccades. Journal of Neurophysiology, 80(5), 2274–2294. https://doi.org/10.1152/jn.1998.80.5.2274

44. Klier, E. M., Wang, H., & Crawford, J. D. (2001). The superior colliculus encodes gaze commands in retinal coordinates. Nature Neuroscience, 4(6), 627–632. https://doi.org/10.1038/88450

45. Klinghammer, M., Blohm, G., & Fiehler, K. (2015). Contextual factors determine the use of allocentric information for reaching in a naturalistic scene. Journal of Vision, 15(13), 1–13. https://doi.org/10.1167/15.13.24.doi

46. Klinghammer, M., Blohm, G., & Fiehler, K. (2017). Scene configuration and object reliability affect the use of allocentric information for memory-guided reaching. Frontiers in Neuroscience, 11(APR), 1– 13. https://doi.org/10.3389/fnins.2017.00204

47. Knight, T. A. (2012). Contribution of the frontal eye field to gaze shifts in the head-unrestrained rhesus monkey: Neuronal activity. Journal of Neuroscience, 225, 213–236. https://doi.org/10.1016/j.neuroscience.2012.08.050

48. Knight, T. A., & Fuchs, A. F. (2007). Contribution of the frontal eye field to gaze shifts in the head-unrestrained monkey: Effects of microstimulation. Journal of Neurophysiology, 97(1), 618–634. https://doi.org/10.1152/jn.00256.2006

49. Kriegeskorte, N. (2015). Deep Neural Networks : A New Framework for Modeling Biological Vision and Brain Information Processing. The Annual Review of Vision Science, 1, 417–446. https://doi.org/10.1146/annurev-vision-082114-035447

50. Körding, K. P., Beierholm, U., Ma, W. J., Quartz, S., Tenenbaum, J. B., & Shams, L. (2007). Causal inference in multisensory perception. PLoS ONE, 2(9). https://doi.org/10.1371/journal.pone.0000943

51. Lew, T. F., Vul, E. (2015). Ensemble clustering in visual working memory biases location memories and reduces the weber noise of relative positions. Journal of Vision. 15:10.

52. Li, J., Sajad, A., Marino, R., Yan, X., Sun, S., Wang, H., & Crawford, J. D. (2017). Effect of allocentric landmarks on primate gaze behavior in a cue conflict task. Journal of Vision, 17(5). https://doi.org/10.1167/17.5.20

53. Lindsay, G.W. (2021). Convolutional Neural Networks as a Model of the Visual System: Past, Present, and Future. J Cogn Neurosci.33(10): 2017–2031. https://doi.org/10.1162/jocn_a_01544. PMID:32027584

54. Liu, L., Ouyang, W., Wang, X., Fieguth, P., Chen, J., Liu, X., & Pietikäinen, M. (2020). Deep Learning for Generic Object Detection: A Survey. International Journal of Computer Vision, 128(2), 261–318. https://doi.org/10.1007/s11263-019-01247-4

55. Lu Z, & Fiehler K. (2020). Spatial updating of allocentric land-mark information in real-time and memory-guided reaching. Cortex. 125:203–214. https://doi.org/10.1016/j.cortex.2019.12.010

56. Ma, W. J., Beck, J. M., Latham, P. E., & Pouget, A. (2006). Bayesian inference with probabilistic population codes. Nature Neuroscience, 9(11), 1432–1438. https://doi.org/10.1038/nn1790

57. Mikula, L., Gaveau, V., Pisella, L., Khan, Z. A., Blohm, G. (2018). Learned rather than online relative weighting of visual-proprioceptive sensory cues. J Neuro-physiol 119: 1981–1992. https://doi.org/10.1152/jn.00338.2017

58. Monga, V., Li, Y., Eldar, Y. C. (2020). Algorithm unrolling: Interpretable, efficient deep learning for signal and image processing, IEEE Signal Processing Magazine.

59. Naka, K. I., & Rushton, W. A. H. (1966). An attempt to analyse colour reception by electrophysiology. The Journal of Physiology, 185(3), 556–586. https://doi.org/10.1113/jphysiol.1966.sp008002

60. Neggers, S. F. W., Schölvinck, M. L., & van der Lubbe, R. H. J. 2005. Quantifying the interactions between allo- and egocentric representations of space. Acta Psychol (Amst). 118:25–45.

61. Neggers, S. F. W., Van der Lubbe, R. H. J., Ramsey, N. F., & Postma, A. (2006). Interactions between ego- and allocentric neuronal representations of space. NeuroImage, 31(1), 320–331. https://doi.org/10.1016/j.neuroimage.2005.12.028

62. Orhan, A. E., & Ma, W. J. (2017). Efficient probabilistic inference in generic neural networks trained with non-probabilistic feedback. Nature Communications, 8(1). https://doi.org/10.1038/s41467-017-00181-8

63. Pitkow, X., & Angelaki, D. E. (2014). Perspective How the Brain Might Work : Statistics Flowing in Redundant Population Codes. Perspective, 1–9. https://doi.org/10.1016/j.neuron.2017.05.028

64. Pouget, A., & Snyder, L. H. (2000). Computational approaches to sensorimotor transformations. Nature Neuroscience, 3(11s), 1192–1198. https://doi.org/10.1038/81469

65. Rolls, E. T. (2020). Spatial coordinate transforms linking the allocentric hippocampal and egocentric parietal primate brain systems for memory, action in space, and navigation. Hippocampus, 30(4), 332–353. https://doi.org/10.1002/hipo.23171

66. Rajalingham, R., Issa, E. B., Bashivan, P., Kar, K., Schmidt, K., & DiCarlo, J. J. (2018). Large-scale, high-resolution comparison of the core visual object recognition behavior of humans, monkeys, and state-of-the-art deep artificial neural networks. Journal of Neuroscience, 38(33), 7255–7269. https://doi.org/10.1523/JNEUROSCI.0388-18.2018

67. Sadeh, M., Sajad, A., Wang, H., Yan, X., & Crawford, J. D. (2015). Spatial transformations between superior colliculus visual and motor response fields during head-unrestrained gaze shifts. European Journal of Neuroscience, 42(11), 2934–2951. https://doi.org/10.1111/ejn.13093

68. Sajad, A., Sadeh, M., Yan, X., Wang, H., & Crawford, J. D. (2015). Visual–Motor Transformations Within Frontal Eye Fields During Head-Unrestrained Gaze Shifts in the Monkey. Cerebral Cortex, 10(1), 1– 8.

69. Sajad, A., Sadeh, M., Yan, X., Wang, H., & Crawford, J. D. (2016). Transition from target to gaze coding in primate frontal eye field during memory delay and memory-motor transformation. ENeuro, 3(2), 82. https://doi.org/10.1523/ENEURO.0040-16.2016

70. Salinas, E., & Abbott, L. F. (1994). Vector reconstruction from firing rates. Journal of Computational Neuroscience, 1(1–2), 89–107. https://doi.org/10.1007/BF00962720

71. Salinas, E., & Abbott, L. F. (1995). Transfer of coded information from sensory to motor networks. Journal of Neuroscience, 15(10), 6461–6474. https://doi.org/10.1523/jneurosci.15-10-06461.1995

72. Schenk, T. (2006). An allocentric rather than perceptual deficit in patient D.F. Nature Neuroscience, 9(11), 1369–1370. https://doi.org/10.1038/nn1784

73. Schrimpf, M., Kubilius, J., Hong, H., Majaj, N. J., Rajalingham, R., Issa, E. B., Kar, K., Bashivan, P., Prescott-Roy, J., Schmidt, K., Yamins, D. L. K., & DiCarlo, J. J. (2018). Brain-Score: Which Artificial Neural Network for Object Recognition is most Brain-Like? BioRxiv. https://doi.org/10.1101/407007

74. Scott, S. H. (2001). Vision to action: New insights from a flip of the wrist. Nature Neuroscience, 4(10), 969–970. https://doi.org/10.1038/nn1001-969

75. Serre, T., Kouh, M., Cadieu, C., Knoblich, U., Kreiman, G., Poggio, T. (2005). A Theory of Object Recognition: Computations and Circuits in the Feedforward Path of the Ventral Stream in Primate Visual Cortex. Technical report.

76. Smith, M. A., & Crawford, J. D. (2005). Distributed population mechanism for the 3-D oculomotor reference frame transformation. Journal of Neurophysiology, 93(3), 1742–1761. https://doi.org/10.1152/jn.00306.2004

77. Soechting, J. F., & Flanders, M. (1992). Moving in three dimensional space: frames of reference, vectors, and coordinate systems. Annual Review of Neuroscience, 167–191.

78. Sommer, M. A., & Wurtz, R. H. (2006). Influence of the thalamus on spatial visual processing in frontal cortex. Nature, (444), 347–377.

79. Thaler, L., & Goodale, M. A. (2011). The role of online visual feedback for the control of target-directed and allocentric hand movements. Journal of Neurophysiology, 105(2), 846–859. https://doi.org/10.1152/jn.00743.2010

80. Tweed, D., Cadera, W., & Vilis, T. (1990) Computing three-dimensional eye position quaternions and eye velocity from search coil signals. Vis. Res. 30: 97–110. https://doi.org/10.1016/0042-6989(90)90130-D

81. Wang, X., Zhang, M., Cohen, I. S., & Goldberg, M. E. (2007). The proprioceptive representation of eye position in monkey primary somatosensory cortex. Nature Neuroscience, 10(5), 640–646. https://doi.org/10.1038/nn1878

82. Webb, B. S., Ledgeway, T., & Rocchi, F. (2011). Neural computations governing spatiotemporal pooling of visual motion signals in humans. Journal of Neuroscience, 31(13), 4917–4925. https://doi.org/10.1523/JNEUROSCI.6185-10.2011

83. Xing, J., & Andersen, R. A. (2000). Models of the posterior parietal cortex which perform multimodal integration and represent space in several coordinate frames. Journal of Cognitive Neuroscience, 12(4), 601–614. https://doi.org/10.1162/089892900562363

84. Zhao, H., & Wildes, R. P. (2021). Interpretable deep feature propagation for early action recognition, arXiv:2107.05122v1.

85. Zipser, D., & Andersen, R. A. (1988). A back-propagation programmed network that simulates response properties of a subset of posterior parietal neurons. Nature, 336, 403–405. https://doi.org/10.1038/334242a0

