## Supplementary material for "Integration of allocentric and egocentric visual information in a convolutional / multilayer perceptron network model of goal-directed gaze shifts": Supplemntal figures

### Supplementary figures

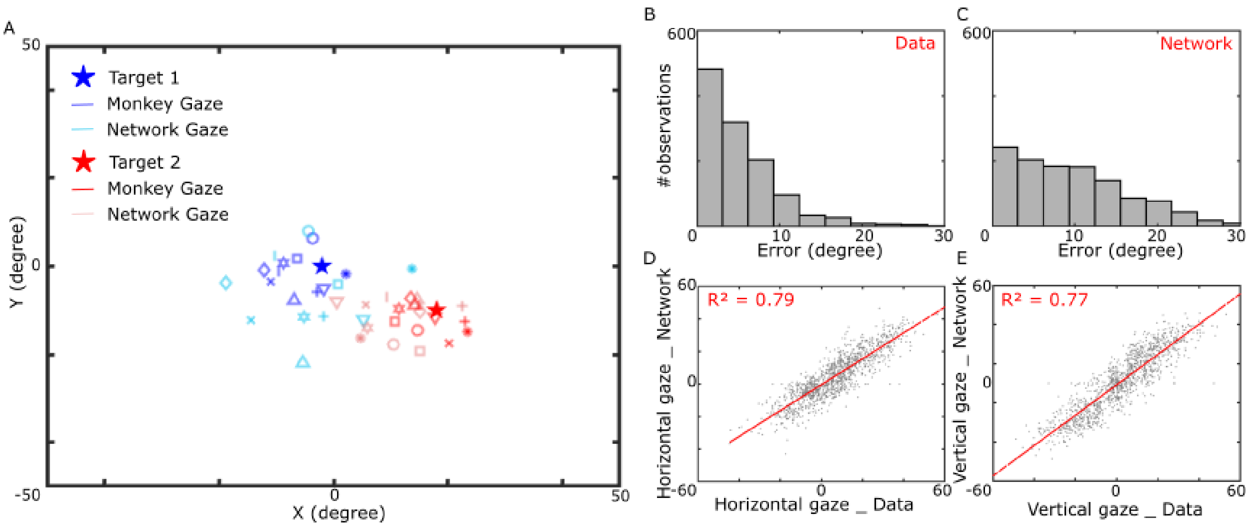


***Figure S1. Network performance for Monkey #2.*** *A) Gaze endpoint errors observed from monkey data. The errors are mainly bellow 8-12°. B) Gaze endpoint errors generated by our network. The majority of errors are below 8-12°, similar to the monkey data. C-E) Regression analysis of our network gaze endpoints and the observed data. Our network explained approximately 80% of the data's variability for both horizontal and vertical directions.*

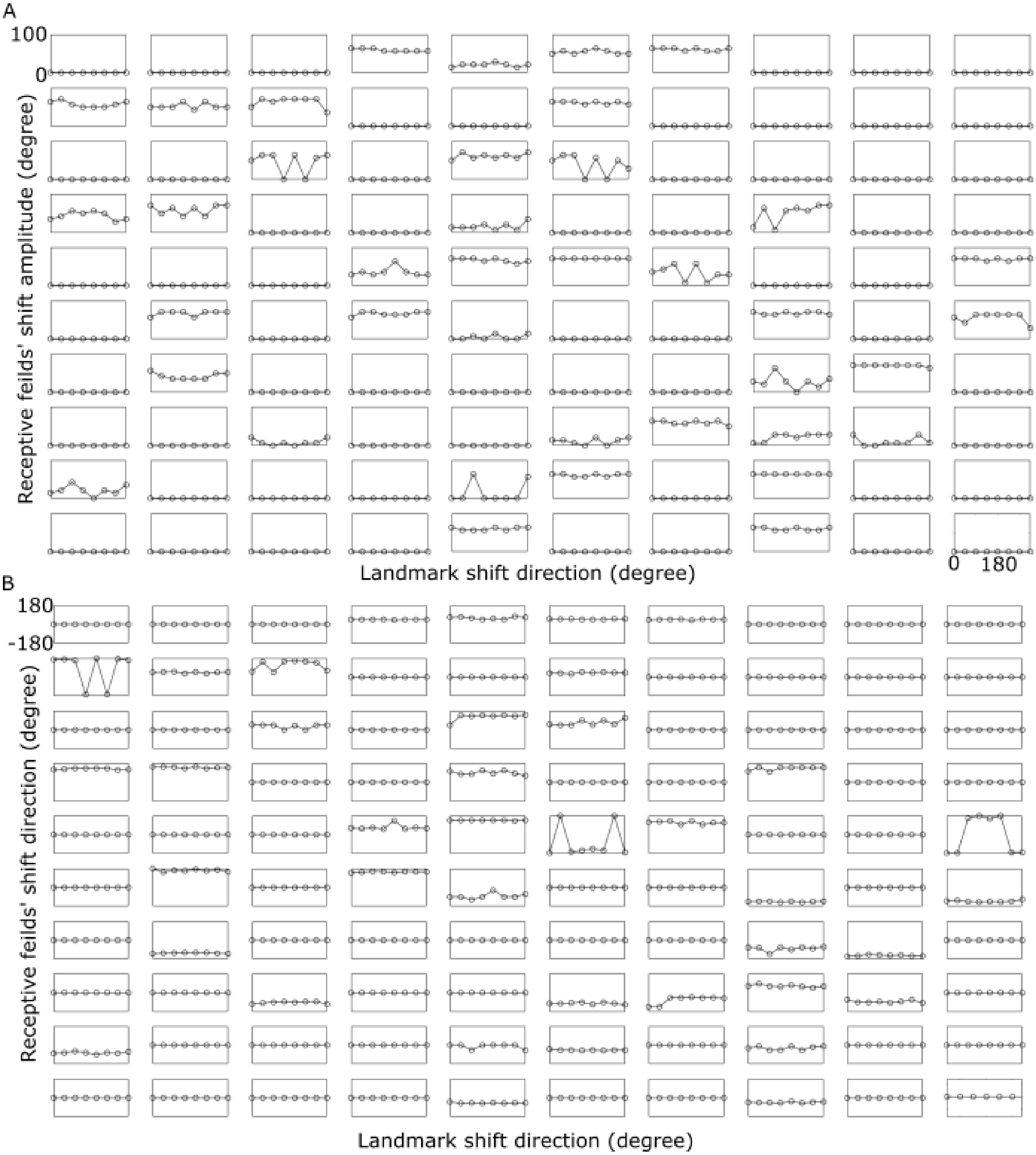


***Figure S2. Quantifying receptive field shifts caused by landmark shifts.*** *Each box represents a unit in the motor population unit. A: amplitude (0-100°) of response field shift plotted as a function of landmark shift direction (-180-180°). B: direction of response field shift (-180-180°) plotted as a function of landmark shift direction (-180-180°). To obtain these data, receptive fields' Peak of activity for each landmark shift was subtracted from the no landmark shift condition. As can be seen, different neurons coded different landmark shift amounts and directions.*

#### Supplementary Method: Determining intrinsic coordinate frames

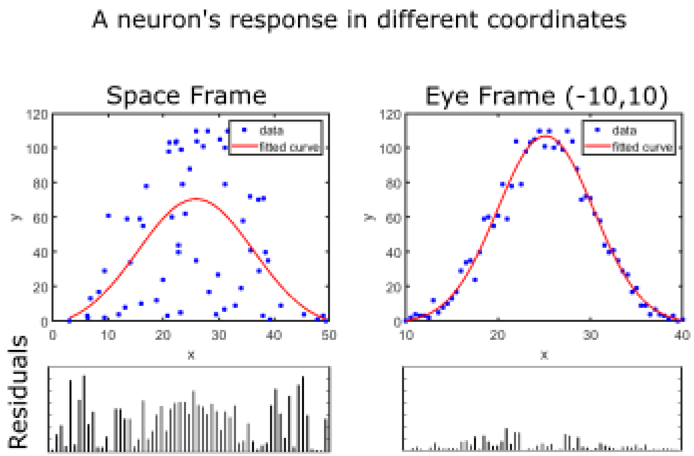


***Figure S3. Determining a neuron's intrinsic coordinate frames.*** *A simulated neuron's response with a Gaussian receptive field coding information in eye coordinates is plotted (right column). The initial eye position varied between [-10,10]. When the neuron is plotted and fit in the wrong space coordinate frame, (left columns), the fit curve fails to closely follow the neuron's response and returns high residuals (lower box).*

To illustrate our fitting methodology, we provided a simple pair of synthetic examples. For this example, we generated responses of a neuron with a Gaussian response field that codes the information in eye coordinates. Eye positions were randomly selected from a uniform distribution in the interval [-10,10]. Figure S2 illustrates our simulated neuron's response (blue dots) in two different spatial coordinates, Space-centered and Eye-centered frame of reference. The red line represents the result of the curve fit for this simulated neuron. As can be seen in the left column, since the neuron response is plotted and fit in the incorrect coordinate frame, the fit curve fails to follow the neuron's response closely, which results in high errors (residuals, lower box). Second, to emphasize further the importance of the intrinsic coordinate frames, we ran a similar simulation in the correct coordinate frames, eye frames. However, when the neuron's response is plotted and fitted in eye coordinates, the curve follows the neuron's response closely (right column), which results in lower errors (residuals, lower box).
